# Mouse parabrachial neurons signal a relationship between bitter taste and nociceptive stimuli

**DOI:** 10.1101/383661

**Authors:** Jinrong Li, Christian H. Lemon

## Abstract

Taste and somatosensation both partly mediate protective behaviors. Bitter taste guides avoidance of ingestion of toxins while pain sensations, such as noxious heat, signal adverse conditions to ward off harm. Although brain pathways for taste and somatosensation are typically studied independently, prior data suggest they intersect, potentially reflecting their common protective role. To investigate this, we applied electrophysiologic and optogenetic techniques in anesthetized mice of both sexes to evaluate relationships between oral somatosensory and taste activity in the parabrachial nucleus (PbN), implicated for roles in gustation and pain. Spikes were recorded from taste-active PbN neurons tested with oral delivery of thermal and chemesthetic stimuli, including agonists of nocisensitive transient receptor potential (TRP) ion channels on somatosensory fibers. Gustatory neurons were also tested to follow electrical pulse stimulation of an oral somatosensory region of the spinal trigeminal subnucleus caudalis (Vc), which projects to the PbN. Neurons composed classic taste groups, including sodium, electrolyte, appetitive, or bitter oriented cells. Across groups, most neurons spiked to Vc pulse stimulation, implying trigeminal projections reach PbN gustatory neurons. Among such cells, agonists of nocisensitive TRP channels, including mustard oil, capsaicin, and noxious heat, were discovered to predominantly activate PbN bitter taste neurons tuned to the bitters quinine and cycloheximide. Such neurons populated the lateral PbN. Further, PbN bitter taste neurons showed suppressed oral nociceptive activity during optogenetic-assisted inhibition of the Vc, implying convergent trigeminal input contributed to such activity. Our results imply a novel role for PbN gustatory cells in crossmodal signaling related to protection.

## Introduction

The mouth is richly innervated by the sensory receptors and endings of neural circuits for taste and somatosensation. Although mediating different sensory qualities, both of these modalities share the function of protective signaling against conditions associated with physiological distress. Trigeminal neurons, the primary mediators of craniofacial somatosensation, can respond to stimuli associated with tissue injury, including oral delivery of noxious heat (> 43°C, Zotterman, 1936; Yarmolinsky et al., 2016) and chemical agonists of transient receptor potential (TRP) ion channels involved with pain sensitivity, such as the capsaicin receptor TRP vanilloid 1 (TRPV1; Carstens et al., 1998; Caterina et al., 2000). Trigeminal neurons can also fire to oral presence of ingesta that would challenge homeostasis, such as highly concentrated salts (Carstens et al., 1998). Relatedly, receptor cells and neurons of the gustatory system mediate signals linked to protection from consumption of toxins (Scott and Mark, 1987), which can taste sour or bitter to humans (Reed and Knaapila, 2010). Moreover, the central neural processing of innately non-aversive taste stimuli, such as sweeteners, can adaptively change to reflect pairing these tastes with negative ingestive consequences (Chang and Scott, 1984; Grossman et al., 2008).

In addition to overlapping function, neural pathways for taste and somatosensation physically overlap and intersect at multiple levels of the nervous system, including early stages of the neuraxis. The lingual branch of the trigeminal nerve partly innervates tongue tissue containing taste buds innervated by the chorda tympani (CT) nerve, a major taste-active facial nerve afferent (Abe et al., 2005; Dhaka et al., 2008). Excitation of the lingual nerve can modulate gustatory firing in neurons of the CT and also the nucleus of the solitary tract (NTS), the first central relay for taste, potentially through axonal reflex release of peripheral neuromodulators (Wang et al., 1995; Simons et al., 2003a). Moreover, although most trigeminal fibers enter the brain to project to the hindbrain trigeminal sensory complex (Marfurt and Rajchert, 1991), some reach CT-supplied neurons in the NTS (Felizardo et al., 2009). Accordingly, electrical stimulation of mandibular trigeminal afferents can excite and modify sensory firing in taste-active NTS neurons (Boucher et al., 2003; Braud et al., 2012; Li and Lemon, 2015a).

While the function of physiological ties between taste and somatosensory circuits is unknown, such phenomena could reflect the common role of both systems in protective coding. In rodents, data relevant to this question stem from studies on the parabrachial nucleus (PbN) of the pons, which is implicated for affective processing in gustation (Yamamoto et al., 1994) and pain (Gauriau and Bernard, 2002). Central tract tracing work revealed the dorsal spinal trigeminal subnucleus caudalis (Vc), involved in part with oral nociceptive processing (Carstens et al., 1998), projects to regions of the PbN containing gustatory-responsive neurons, implying PbN circuits support convergence of trigeminal sensory signals with taste (Cechetto et al., 1985). Further, independent functional studies on gustation or pain identified that oral delivery of bitter tastants (Yamamoto et al., 1994; Halsell and Travers, 1997; Geran and Travers, 2009) and noxious stimulation, including heat or pinch, of trigeminal receptive fields (Bernard and Besson, 1990; Campos et al., 2018) excite neurons in common PbN subregions, including the external lateral and external medial PbN subnuclei. The above data open the possibility that aversive somatosensory and gustatory signals could activate the same PbN neurons.

To test this idea, we made in vivo electrophysiological recordings from gustatory-active PbN neurons stimulated with oral thermal and chemesthetic stimuli associated with somatosensory and pain transmission.

Electrical and optogenetic perturbation directed to Vc axonal projections to the PbN evaluated connectivity of taste-active PbN cells with upstream trigeminal circuits. Our results provide evidence that ascending trigeminal projections reach multiple types of taste-identified neurons in the PbN and specifically imbue a subpopulation of bitter taste-responsive cells with sensitivity to oral nociceptive stimuli. Such “gustatory” neurons appear to convey crossmodal signals relevant to protective coding and to mediate functions beyond unimodal taste.

## Materials and Methods

### Animals and preparation

These studies used adult mice of both sexes from the C57BL/6J (B6) line (stock #000664, Jackson Labs, Bar Harbor, ME) and mice derived from a cross between B6 mice and an optogenetic inhibitory mouse line, discussed below. All mice were naïve to experimentation and were housed in a vivarium that maintained a 12:12h light:dark cycle and an air temperature of around 20°C. Food and water were available ad libitum.

In brief, the procedures described below allowed for acute recording of extracellular spikes from taste-active PbN neurons in anesthetized mice during controlled stimulation of the oral cavity with gustatory and somatosensory stimuli. We also tested whether the recorded PbN units fired spikes in response to weak electrical pulses delivered through the tip of a stimulation electrode located in the orosensory Vc, thus evaluating functional coupling of central trigeminal circuits with gustatory cells. Finally, a fiber optic probe was positioned alongside the Vc stimulation electrode when recording from optogenetic inhibitory mice to enable tests on how reversible light-induced suppression of trigeminal circuits impacted somatosensory firing in taste-active PbN cells.

Mice were prepared for neurophysiologic and optogenetic experiments with guidance from the University of Oklahoma Institutional Animal Care and Use Committee, which reviewed and approved our procedures, and the National Institutes of Health. To ease injection of a short-acting anesthetic/analgesic agent needed for initial preparation, mice were briefly placed in a small plexiglass box infused with 4-5% isoflurane in oxygen until breathing was regularly slow and body movement was absent. Mice were then inducted using a mixture of ketamine (100 mg/kg, i.p.) and xylazine (10 mg/kg, i.p.). Atropine (24 μg/kg, i.p.) was also administered to reduce bronchial secretions. Once anesthetized as evidenced by absence of a reflex response, a tracheostomy tube (PE 60, length = 25 mm) was inserted to allow breathing during oral stimulation with liquids. To facilitate access to the oral cavity as discussed below, a small suture was placed through the superficial rostroventral tongue and the mandibular incisors were partly trimmed using rongeurs. Mice were then secured in a stereotaxic instrument with ear bars (Model 930, David Kopf Instruments, Tujunga, CA). For long-duration anesthesia needed for recording studies, the open end of the tracheostomy tube was coupled to concentric pressure-vacuum gas exchange tubing (Wilson and Lemon, 2014) through which mice freely inhaled 0.8 to 1.2% isoflurane in oxygen. Under this system, mice respired anesthetic gas without the aid of an external ventilator. Body temperature was kept at around 37.4°C by a feedback-controlled heating pad.

Anesthesia was monitored and confirmed by absence of reflex to heavy pinch applied to a hind paw. Under anesthesia, the scalp was incised along the midline of the skull to expose bregma and lambda, which were brought into the same dorsoventral plane through stereotaxic adjustments. The mandible was gently deflected downward and held in position by a small thread of surgical silk that passed caudal of the incisors and was fixed to the stereotaxic device. The tongue was extended from the mouth by light pressure on the rostroventral suture. To allow electrode access to the PbN, a unilateral craniotomy removed a small portion of the interparietal bone and exposed the surface of inferior colliculus and the parts of the cerebellum. To allow electrode and fiber optic access to the Vc, the caudal zone of the occipital bone ipsilateral to the targeted PbN was trimmed. Dura mater overlying the inferior colliculus and targeted region of the medulla was retracted to facilitate electrode passage into brain tissue.

A concentric bipolar stimulation electrode (CEA-200, Microprobes, Gaithersburg, MD) was targeted to the Vc using a micromanipulator (SM-15R, Narishige International, Amityville, NY) attached to the stereotaxic device. For optogenetic studies involving light-induced neural suppression also directed to the Vc, the cut and prepared end of a 200 μm diameter fiber optic cable (0.39 numerical aperture, ThorLabs, Newton, NJ)was coupled alongside the Vc stimulation electrode, with the tip of the fiber optic positioned just behind the electrode tip to enable light delivery to the brain area penetrated by the electrode. Coordinates for the stimulation electrode targeted the oral thermal/sensory Vc in mice, as based on our prior work (Lemon et al., 2016): approximately 8.0 to 8.2 mm caudal of bregma, approximately 1.8 to 2.2 mm lateral of the midline, and superficial depth just below the medullary surface. The stimulation electrode approached the Vc at an approximate 15° angle in the sagittal plane to allow space for PbN access. Final positioning of the stimulating electrode was confirmed by electrophysiological monitoring for the first signs of multiunit activity to oral presentation of cool water following 35°C adaptation, as below. Once in position, the core and outer leads of the stimulating electrode were electrically connected to, respectively, the positive and negative poles of a constant current stimulus isolation unit (PSIU6X, Grass Technologies) coupled to a programmable square pulse stimulator (S88X, Grass Technologies).

Following positioning of the Vc stimulation electrode, the tip of the recording electrode (custom 2 to 5 MΩ single-channel tungsten probe, FHC Inc., Bowdoinham, ME) was targeted to the PbN using stereotaxic coordinates: 4.7 to 5.0 mm caudal of bregma, 1.2 to 1.5 mm lateral of the midline, and 2.3 to 3.0 mm below the brain surface. The dorsoventral position of this probe was stepped by an electronic micropositioner (Model 2660, David Kopf Instruments). While lowering the electrode, taste-sensitive neurons were sought out and identified through monitoring for excitatory single-unit activity to 300 mM NaCl, which interspersed continued oral delivery of 35°C water as below. Electrophysiological activity was AC amplified (P511 with high-impedance probe, Grass Technologies), bandpass filtered (around 0.3 to 10 kHz), and monitored on an oscilloscope and loudspeaker. Well-isolated single-neuron spikes were sampled (25 kHz) under a template-matching algorithm (1401 interface and Spike2 software [version 9], CED, Cambridge, England). Time stamps for spikes were captured with a precision of 0.1 ms. Data files for individual trials were downloaded to storage media while recording and analyses were performed offline.

### Electrical stimulation of the Vc

To investigate their potential upstream coupling with trigeminal circuits, taste-sensitive PbN neurons were tested for spike activation following delivery of a train of electrical pulses to the ipsilateral Vc at 1 Hz (pulse amplitude ≈ 150 μA; pulse width = 500 μs). Each pulse was considered to mark a trial, with the 100 ms period prior to the pulse referenced as baseline/random firing. Over many trials, spikes that arose within a 200 ms post-pulse window were statistically compared to baseline activity to determine whether the unit reliably spiked in response to electrical stimulation of the Vc, as below. Note this procedure did not intend to reveal the number of synapses involved but evaluated electrophysiologic coupling between the stimulated brain site and recorded PbN neurons.

### Oral stimuli

PbN neurons identified to respond to taste input were tested for sensitivity to oral presence of solutions of diverse gustatory chemicals, thermal stimuli in the form of temperature-adjusted volumes of purified water, and solutions of chemesthetic stimuli including capsaicin. Capsaicin was applied to lingual tissue in a unique manner, discussed below. All other stimuli were stored in airtight glass bottles and thermally controlled by submersion in microprocessor-regulated warming and recirculated cooling/heating water baths. Temperature-controlled stimulus solutions were flowed into the mouse oral cavity on individual trials using a custom apparatus operated and timed by the data acquisition system, as in our prior studies (Wilson and Lemon, 2014; Lemon et al., 2016). Briefly, on a given trial, a 3-way valve seamlessly switched oral solution flow from a 35°C (i.e., near physiological temperature) purified water adaptation rinse to a stimulus solution delivered for a brief period, following which flow returned to the 35°C adaptation rinse. The adaptation rinse continued in between trials to facilitate thermal adaptation of oral tissue to physiological value. Inter-trial intervals were around 2 m. Solutions flowed into the mouth at a rate of around 1.5 ml/s and bathed anterior and posterior oral fields (i.e., whole-mouth stimulation; Wilson et al., 2012).

Stimuli were loaded into the delivery system by funnel, with effort made to take about the same amount of time to prepare and start each trial for thermal consistency. As in our prior work (e.g., Lemon et al., 2016), solution flow temperature was continuously monitored at the moment of oral entry using a small thermocouple probe (IT-1 E, time constant = 0.005 s, Physitemp Instruments, Inc., Clifton, NJ) connected to a digital thermometer (BAT-12, precision = 0.1 °C, Physitemp Instruments, Inc.). The analog output of the thermometer was sampled (1 kHz) by the data acquisition system to give real-time measurement of the temperature of oral solution flow alongside cellular spiking data. Stimulus temperatures were quantified as mean oral temperature across trials during the last 3 s of stimulus delivery. In between trials, the funnel and all stimulus passages of the delivery system were rinsed with 35°C purified water.

Taste, thermal, and chemesthetic stimuli were respectively tested in 3 separate blocks. Taste stimuli were 500 mM sucrose (SUC), a mixture of 100 mM monopotassium glutamate and 10 mM inosine 5’-monophophate (UMA), 100 mM NaCl (NAC), 10 mM citric acid (CIT), 10 mM quinine-HCl (QUI) and 0.1 mM cycloheximide (CHX). These stimuli are associated with the sweet (SUC), umami (UMA), salty (NAC), sour (CIT), and bitter (QUI and CHX) human taste categories. Stimulus concentrations were selected from prior gustatory electrophysiology studies conducted in mouse brain stem (Wilson et al., 2012; Tokita and Boughter, 2016). Notably, the bitters QUI and CHX induce strong oral sensory aversion in naïve B6 mice (Boughter et al., 2005; Ellingson et al., 2009). Taste chemicals were of high purity (purchased from Sigma, St. Louis, MO) and dissolved in purified water. Taste solutions were delivered for 5 s at an oral temperature of 28°C. The ordering of taste chemical trials was generally randomized, without replacement, for each cell.

Thermal stimuli were 5 s presentations of purified water adjusted to each of five temperatures, as measured inside the mouth: 14°, 21°, 28°, 35° and 48°C. These temperatures included noxious heat (48°C, abbreviated as HEAT) and spanned broad range of distinct cooling and warming values that can excite oral trigeminal neurons (e.g., Carstens et al., 1998; Lemon et al., 2016). Note temperatures below physiological value (35°C) were considered “cool”. Thermal trial order was randomized.

Finally, the chemesthetic stimuli tested were 28°C 1.28 mM L-menthol (MENT), 35°C 1 mM allyl isothiocyanate (mustard oil, abbreviated AITC), and room temperature 1 mM capsaicin (CAP). Capsaicin is an agonist of TRPV1 expressed by somatosensory neurons (Caterina et al., 1997); Mustard oil stimulates TRP ankyrin 1 (TRPA1; Jordt et al., 2004) and partly TRPV1 (Everaerts et al., 2011); Menthol is associated with activation of TRP melastatin 8 (TRPM8; McKemy et al., 2002). Wild-type and B6 mice show strong ingestive and oral sensory avoidance of CAP and AITC (Ellingson et al., 2009; Everaerts et al., 2011). In B6 mice, MENT can electrophysiologically excite trigeminal ganglion neurons when delivered orally but cause only relatively mild avoidance in brief-access drinking tests (our unpublished observations). MENT and AITC were dissolved in purified water. CAP was dissolved in a vehicle of 1.5% ethanol/1.5% Tween 80 in purified water; PbN neurons were also tested with the CAP vehicle on a separate trial. Solutions of MENT and AITC were presented orally for 10 s using our stimulus delivery system. For CAP and CAP vehicle trials, a small volume (estimated 150 μL) of the solution was applied to a cotton-tipped applicator that was lightly dabbed onto the exposed rostral-ipsilateral tongue for 20 s; during this period, delivery of the oral adaptation rinse was paused. Postmortem inspection of a mouse tongue where thionin dye was applied in the same manner revealed stimuli dabbed by cotton swab were restricted to tongue regions rostral of the median eminence and did not reach caudal foliate or circumvallate taste papillae. While the order of MENT and AITC trials varied across cells, the CAP trial was always tested after the CAP vehicle, at the end of the chemesthetic stimulus block, as CAP delivery induced lingering responses and effects (see also Carstens et al., 1998).

If the PbN neuron remained isolated and displayed baseline firing approximately 20 m after completion of the CAP trial, it was tested with a concentration series of NaCl that included 10, 30, 100, 300, and 1000 mM solutions presented at 28°C. This series included high concentrations of NaCl (300 and 1000 mM) that can elicit intake avoidance in B6 mice (Ninomiya et al., 1989; Eylam and Spector, 2002). The testing order for NaCl concentrations was randomized for each cell.

### Optogenetic-assisted suppression of Vc oral activity during PbN recordings

Optogenetics was used to reversibly inhibit Vc circuitry during recordings from Vc-coupled PbN neurons to evaluate how temporary removal of ascending trigeminal input affected PbN firing over multiple trials to the bitter tastant QUI and nociceptive stimulus HEAT. For these tests, HEAT was used as a nociceptive stimulus because it induced activity in PbN neurons that readily reverted to baseline on stimulus cessation, which avoided lasting neural excitation and carryover effects that could result from application of a chemonociceptive stimulus, such as CAP (e.g., Figure 4). We used a silencing by excitation strategy (Wiegert et al., 2017) where the natural inhibitory circuitry of the Vc was acutely engaged by light to suppress activity in an oral sensory-verified region of this nucleus. The Vc is heavily populated with GABAergic cells (Ginestal and Matute, 1993) that do not project to the PbN (Haring et al., 1990) and likely modulate intra-Vc processing and Vc outputs. The location of oral sensory neurons in the superficial dorsal Vc (Carstens et al., 1998; Lemon et al., 2016) eased light delivery to this circuit.

Mice used for optogenetic studies were bred from a cross between wild-type B6 mice and hemizygous VGAT-ChR2-EYFP BAC line 8 mice (Jackson Labs, Bar Harbor, ME: 014548; Zhao et al., 2011). The latter express the excitatory opsin Channelrhodopsin-2 (ChR2) and an enhanced yellow fluorescent protein (EYFP) tag in neurons containing the vesicular γ-aminobutyric acid (GABA) transporter (VGAT). VGAT is found in GABAergic cells implicated for inhibition (Sagne et al., 1997). VGAT-ChR2-EYFP mice afford reversible photoinhibition of small volumes of neural tissue through selective activation of local inhibitory interneurons with blue light (Guo et al., 2014; Sofroniew et al., 2015; Wiegert et al., 2017).

The offspring of our cross were identified as optogenetic-excitable (VGAT-ChR2 mice) or optogenetic-negative by the presence or absence, respectively, on post-natal day 0 to 2 of trans-dermal EYFP illumination in VGAT-positive spinal cord interneurons (MZ10 F stereomicroscope with EYFP fluorescence, Leica Microsystems Inc., Buffalo Grove, IL). VGAT-ChR2 mice were also confirmed as optogenetic-positive during setup for electrophysiological testing by monitoring for change in Vc multiunit activity following blue laser pulse stimulation of the Vc, as below. For some of these mice, EYFP fluorescence of brain tissue was assessed and verified following perfusion at the end of PbN unit recordings. EYFP fluorescence was expectedly absent from the brains of all littermate optogenetic-negative mice, which were used as controls for non-specific effects of illumination of brain tissue.

Blue laser light (473 ± 1 nm DPSS laser, OEM Laser Systems, Inc., East Lansing, MI) was delivered to superficial medullary tissues through a fiber optic that was paired, as above, to the electrical stimulation electrode, which targeted oral sensory Vc circuitry. Light irradiance (50 to 70 mW/mm^2^ at fiber tip) enabled activation of ChR2-positive inhibitory circuits influencing oral sensory medullary trigeminal and Vc neurons (e.g., Figure 8A). During testing, stimulus-induced spike activity was repeatedly recorded from a V+ PbN neuron during oral presentation of the stimulus in unison with blue light delivery to the Vc (Vc laser trials) or without laser delivery (control trials). Trials were sequenced and analyzed as below. On Vc laser trials, light delivery was turned on and off at the beginning and end, respectively, of the oral stimulus period using TTL modulation controlled by the data acquisition system. Stimulus periods were shortened to 2 s in optogenetic experiments to accommodate the intent to test single cells with many Vc laser/control trials, as below, and to reduce oral tissue exposure to noxious heat.

### Histology

After data were acquired from the final PbN neuron of the day, mice were overdosed with sodium pentobarbital (≥ 130 mg/kg, i.p.) and weak current (100 μA/1.5 s) was passed through the recording electrode tip to mark its last position. Mice were transcardially perfused with isotonic saline followed by 4% paraformaldehyde/3% sucrose. Brains were removed and stored in a 4% paraformaldehyde/20% sucrose solution. Coronal sections (40 μm) from each brain were mounted onto slides and stained for histological analysis of electrode placement using anatomical landmarks (Franklin and Paxinos, 2008). Note that a majority of, but not all, recording sites were marked or recoverable.

### Experimental design and statistical analysis Electrophysiological recording and stimulation studies on B6 mice

This experiment involved mice of both sexes, using 25 male (mean body weight: 27.0 g ± 2.8 g standard deviation [SD]) and 29 female (22.0 g ± 1.8 g SD) animals. From these mice, 73 taste-active PbN neurons were recorded and tested for electrophysiologic coupling with the Vc. Evidence for or against Vc coupling could not be determined, as below, for 6 cells and 1 additional neuron did not cluster with the major neural groups defined below; these 7 cells were discarded from further analyses. Overall, gustatory response data were analyzed from 66 PbN neurons, albeit only subsets of these cells could be held across the remaining stimulus conditions and trial blocks; cells numbers used for specific analyses are denoted in the results section and figure captions. Sample sizes were comparable to those in the authors’ prior neurophysiological studies on taste and trigeminal processing in mice (Li and Lemon, 2015b; Lemon et al., 2016). Somatosensory-specific PbN neurons were encountered in our recordings but were not included in analyses due to the focus on only taste-active cells.

A 2-way sex × stimulus ANOVA applied to firing to taste stimuli in Hz, calculated as below, by PbN neurons acquired from female (34 cells) and male (32 cells) B6 mice revealed a nonsignificant main effect of sex (*F*_1,64_ = 0.3, *P* = 0.6) and nonsignificant sex × stimulus interaction (*F*_5,320_ = 1.96, *P* = 0.09). Thus, sex was not included as a factor in subsequent analyses. Among analyzed neurons, thirty-four individual and 16 pairs of PbN units were recorded from separate B6 mice. Pairs of neurons were always recorded in sequence, not simultaneously, with the second cell of the pair acquired after repositioning of the PbN recording electrode. Across cells sampled as pairs, no significant correlations were noted between their responses to SUC, UMA, NAC, QUI, or CHX (−0.17 < Spearman’s rank correlation coefficient < 0.23, *P* > 0.4), implying these neurons displayed statistical independence in their firing characteristics.

### Analysis of electrophysiologic coupling between the Vc and PbN neurons

A statistical method was used to evaluate whether PbN neurons reliably generated spikes in response to electrical pulse stimulation directed to the Vc. For Vc stimulation trials (generally 40 were tested) acquired from an individual neuron, time stamps for spikes that followed the electrical pulse (time = 0 ms) were conflated into a single vector. An iterative Poisson-based method (Chase and Young, 2007; Wilson and Lemon, 2014) estimated the probability that a spike density built over sequential spikes in this vector was similar to periods of random firing. A resultant low probability suggested spikes followed Vc stimulation at a rate greater than expected by chance. The time of the evoked spike where this probability became less than 10^−6^ was taken as the neuron’s latency to fire to Vc stimulation. If this latency was less than 100 ms (i.e., within 100 ms of the Vc pulse), the neuron was classified as receiving electrophysiologic input from the Vc (V+ neuron). Otherwise, the cell was considered uncoupled with the Vc (V-neuron).

To ensure that classification of neurons as V− did not result from failure of the electrical stimulation setup, all V− cells included in analyses were acquired from mice where electrical pulse stimulation of the Vc could induce spikes in other PbN neurons. Further, to control for non-specific activation, in two mice the electrical stimulation probe was repositioned to locations medial of the Vc and just across the midline (superficial depth) to determine if pulse stimulation directed to non-Vc tissue could evoke spikes in the recorded PbN neurons. However, in these cases evoked spikes disappeared in PbN units, only to reappear with probe repositioning and pulse stimulation of tissue at the lateral coordinate for the Vc.

### Measurement of stimulus activity

With the exception of CAP trials discussed below, all stimulus responses by PbN cells were quantified as the firing rate during the stimulus period, expressed as spikes per second or Hz, corrected for the baseline firing rate. In these cases, the spike rate during the pre-stimulus period of a trial was subtracted from the firing rate measured during stimulus delivery. For analysis purposes, the stimulus period was considered to lag by 250 ms from the computer-initiated signal for stimulus onset to accommodate transitions between oral adaptation rinse and stimulus flows.

Activity to CAP was also expressed in Hz but was computed differently due in part to the atypical time course of CAP-evoked responses, which could arise with a delay and continue during the post-stimulus rinse period for an extended length of time. Thus, firing on CAP trials was quantified based on a 20 s window that began 10 s prior to the termination of CAP delivery and ended 10 s after this mark. This same window was used to quantify neural firing rates during application of the CAP vehicle, which was the control, or baseline, condition for CAP activity. To compute the corrected CAP response for each cell, the spike rate on the vehicle trial was subtracted from the firing rate computed on the CAP trial.

For some analyses, stimulus responses by individual PbN neurons were converted to standardized firing. For a given neuron, the corrected firing rate computed on each trial was divided by the standard deviation of firing by this cell across all trials considered. This metric was unitless and facilitated analyses of patterns of cross-modal activity with different time course and comparisons among cells that sometimes displayed different general firing rates (Lemon et al., 2016).

### Determination of gustatory neural groups

To group cells by taste sensitivity, hierarchical cluster analysis was applied to a distance matrix for all PbN neurons, where pairwise neural distance was computed as 1 minus the linear correlation between firing in Hz to SUC, UMA, NAC, CIT, QUI, and CHX. Amalgamation proceeded under the average linkage method. Identification of a marked break, or “elbow”, in a line plot cluster distance against amalgamation steps (i.e., a scree plot) revealed the number of neural groups present in the cluster solution (e.g., Lemon et al., 2016). The average firing to taste stimuli by neurons composing each group was used to define gustatory profiles.

### Linear regression analysis of NaCI concentration-response data

For grouped neurons, doubly-logarithmic linear regression was used to test the null hypothesis that their firing in Hz to elevated concentrations of NaCl (100, 300, and 1000 mM) did not change with concentration. A *t*-test with *df* = n − 2 evaluated deviation of the regression line slope from 0. Due to use of logarithmic scales, a value of 1 was added to each response included in this analysis to accommodate a small fraction of just below zero firing rates to NaCl (Wilson and Lemon, 2014).

### Multiple regression analysis

A multiple regression approach was used to explore correlations among activity to nociceptive and bitter taste stimuli in V+ PbN gustatory cells, using nociceptive firing as the dependent measure. Unique correlations between firing to specific nociceptive and bitter stimuli were indexed using semipartial correlation coefficients (*sr*), which can account for redundancies among independent variables (Tabachnick and Fidell, 2001). Regression analyses were performed on standardized firing. To mitigate data heteroscedasticity, weighted regressions were used, where cells with less predictive error for nociceptive firing were, in general, given more weight in the analysis. To compute regression weights, cellular responses to a nociceptive stimulus were first regressed onto firing to bitter stimuli. The absolute values of the residuals from this regression were then regressed onto bitter activity, with, for each neuron, the reciprocal of its resultant squared predicted value taken as its regression weight. Data were inspected for multivariate outliers but none were detected based on calculations of Mahalanobis distance, distributed as *χ*^2^ with *df* = 2 and α = 0.001 (Tabachnick and Fidell, 2001). Regression tolerances did not approach zero, implying collinearity was not an issue.

### Analysis of effects of optogenetic-assisted silencing of the Vc on PbN firing to HEAT and QUI

Four adult male (mean body weight: 34.9 g ± 5.6 g SD) and 9 adult female (27.3 g ± 3.3 g SD) VGAT-ChR2 and littermate control mice were used in these studies. Ten V+ PbN neurons sensitive to HEAT and QUI were acquired from VGAT-ChR2 mice. Six of these neurons yielded response data for both HEAT and QUI on Vc laser and control trials; 2 cells yielded such data for only QUI, and 2 for only HEAT. Data for activity to HEAT and QUI on Vc laser and control trials were acquired from 4 V+ PbN neurons in non-optogenetic littermate mice.

For each stimulus, firing rates in Hz were repeatedly sampled from individual neurons over multiple Vc laser and control trials to account for normal variance in sensory firing. We aimed to record a block of 10 Vc laser and 10 control trials during tests involving one stimulus. An interleaved design was used for trial ordering within a block, where following either an initial control or Vc laser trial (varied across cells), alternating pairs of these trials were presented in sequence. Under this design, testing began with either the control or Vc laser condition and each condition was preceded by itself and the other approximately an equal number of times over the block.

A receiver operating characteristic (ROC) curve (Green and Swets, 1966) quantified the effect of Vc photoinhibition on firing by an individual PbN neuron to HEAT or QUI. The area under each ROC curve (auROC) estimated the probability that random responses to the stimulus sampled under Vc laser and control conditions could be correctly classified by always assuming the larger response was control activity. An auROC that approached 1 reflected very good classification, with auROC = 1 indicating perfect performance. The latter result would arise if stimulus firing was consistently greater on control compared to Vc laser trials, regardless of trial order. On the other hand, an auROC that approached 0.5 indicated near chance discrimination, with auROC = 0.5 arising if the control and Vc laser response distributions completely overlapped. An auROC < 0.5 arose when responses on Vc laser trials were frequently larger than control activity.

A permutation method assessed deviation of an observed auROC value from chance performance. Here, we re-computed 1000 auROC values from the response data for the stimulus where prior to each computation, observed responses were randomly shuffled between Vc laser and control trial labels. This approach resulted in a null distribution of auROC values for the associated stimulus and neuron that reflected random classification performance. An observed auROC that was more extreme than the upper, or lower, 2.5^th^ percentile value of the null distribution was considered to significantly differ from 0.5 (e.g., Veit and Nieder, 2013; Gadziola and Wesson, 2016).

### Other analyses, software, and plotting

In some cases, a bootstrapped procedure estimated the 95% confidence interval of an estimate, denoted as 95% Cl*. In these cases, 1000 bootstrap data resamples were used to calculate the 95% Cl* under the percentile or bias corrected/accelerated percentile method; no interpretative differences were noted between these methods.

Cells that showed significant firing to CAP were identified by visual inspection of spike trains for a robust, lasting spiking response to CAP; this effect was pronounced for a subpopulation of the recorded V+ PbN taste cells (e.g., Figure 4). Significant firing to other oral stimuli was indexed using a statistical algorithm, as based on Wilson and Lemon (2014).

Tests of equal frequencies across conditions were carried out using *χ*^2^ goodness of fit tests. Evoked spike latencies across neural groups were analyzed using a nonparametric Kruskal-Wallis test to accommodate non-normal data. These statistical procedures, along with ANOVA and multiple regression, were carried out using SPSS (version 23.0.0.2, IBM, Armonk, NY). All other calculations and analyses were performed using custom code and built-in functions in MATLAB (release R2018a, MathWorks, Natick, MA).

All data plots and graphs were generated using standard routines and custom code in MATLAB. Raw electrophysiological sweeps and thermal traces were exported in bitmap format from Spike2. Illustrations and final figure configurations were produced using Illustrator (version 22.1, Adobe, San Jose, CA).

## Results

### Analysis of gustatory response data

Cluster analysis applied to gustatory response profiles for 66 taste-active PbN neurons revealed four major cellular groups (Figure 1A). These groups were 1) neurons oriented to NAC (“sodium” cells), 2) neurons with broad tuning across the electrolyte taste stimuli NAC, CIT, and QUI (“electrolyte” cells), 3) neurons oriented toward the preferred taste stimuli SUC and UMA (“appetitive” cells), and 4) neurons that fired strongly and selectively among tastants to the aversive, in unconditioned form, bitter stimuli QUI and CHX (“bitter” cells, Figure 1B). Cells of all types fired to the NaCI concentration series (Figure 2A; Table 1). However, a unique, significant trend was observed for bitter neurons, where more cells significantly fired to high concentrations of NaCI that are innately avoided by B6 mice in behavioral intake tests (300 and 1000 mM, Ninomiya et al., 1989; Eyiam and Spector, 2002) compared to low NaCI concentrations (10 and 30 mM; Table 1). Relatedly, the slope of the concentration-response function for the three highest intensities of NaCI exceeded 0 for bitter neurons (slope = 0.62 [95% Cl lower bound = 0.40; upper = 0.84], *t*_13_ = 6.1, *P* < 0.001) but was indifferent from 0 for all other cellular types (*P* > 0.05); least-squares fits to mean responses are shown in Figure 2B. Thus, increasing the concentration of NaCI to high levels caused the greatest relative increases in responding in bitter neurons, revealing these cells have the highest sensitivity to non-preferred intensities of NaCI.

**Figure 1.**
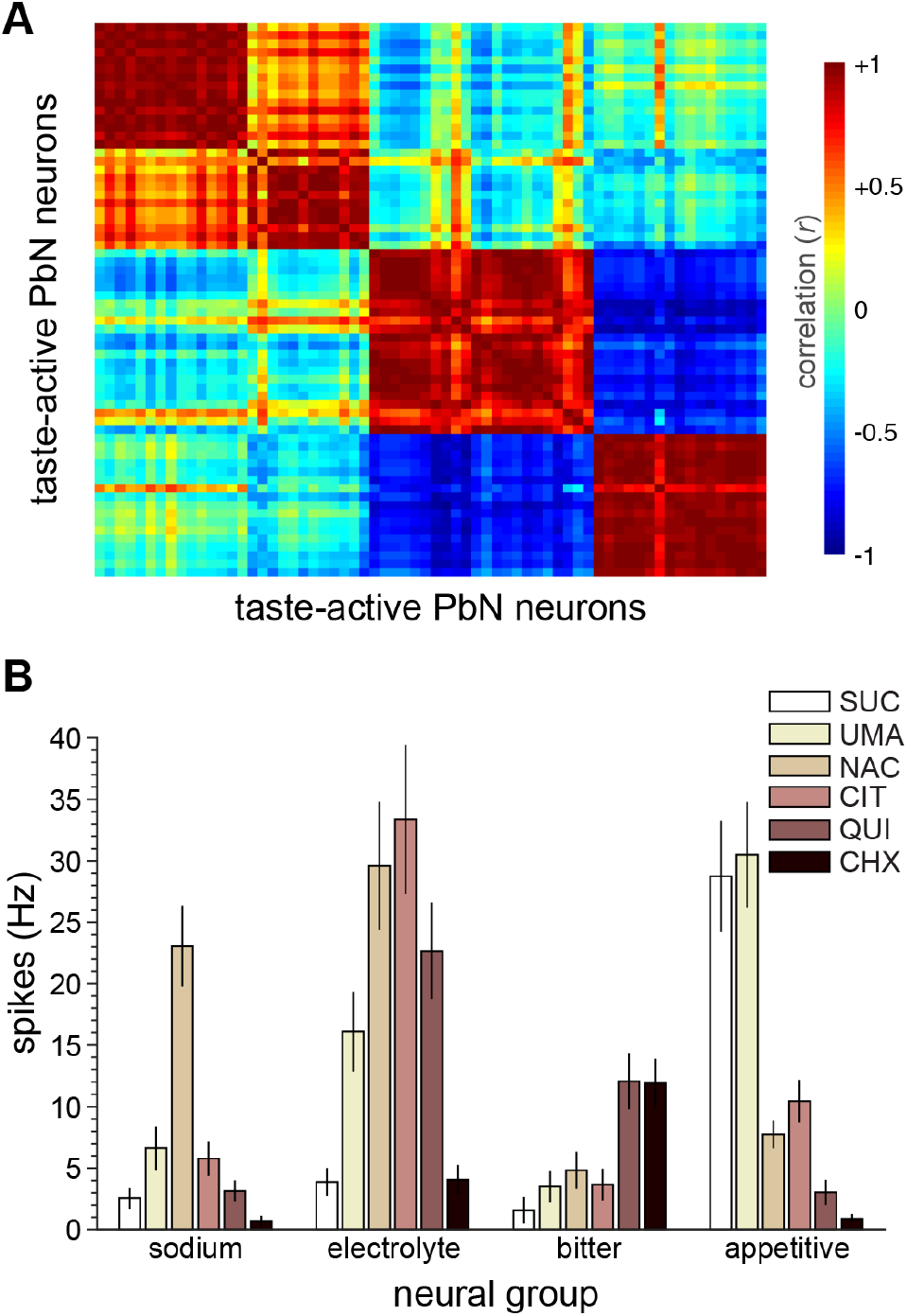
Definition of gustatory neural groups. **A**, Linear correlation matrix for 66 taste-active PbN neurons computed from their firing In Hz to six taste stimuli, as in panel B. Cluster analysis identified four major groups of neurons where cells showed marked positive w¡th¡n-group correlation among their gustatory tuning profiles. Heatmap scale reflects correlation value. **B**, Mean ± s.e.m. firing in Hz by sodium (*n* = 15), electrolyte (*n* = 12), bitter (*n* = 22), and appetitive (*n* = 17) group neurons to six taste stimuli (legend): 500 mM sucrose (SUC), an umami mixture (UMA), 100 mM NaCI (NAC), 10 mM citric acid (CIT), 10 mM quinine (QUI), and 0.1 mM cycloheximide (CHX). All stimuli were thermal-controlled and presented at 28°C following oral adaptation to 35°C.

**Figure 2.**
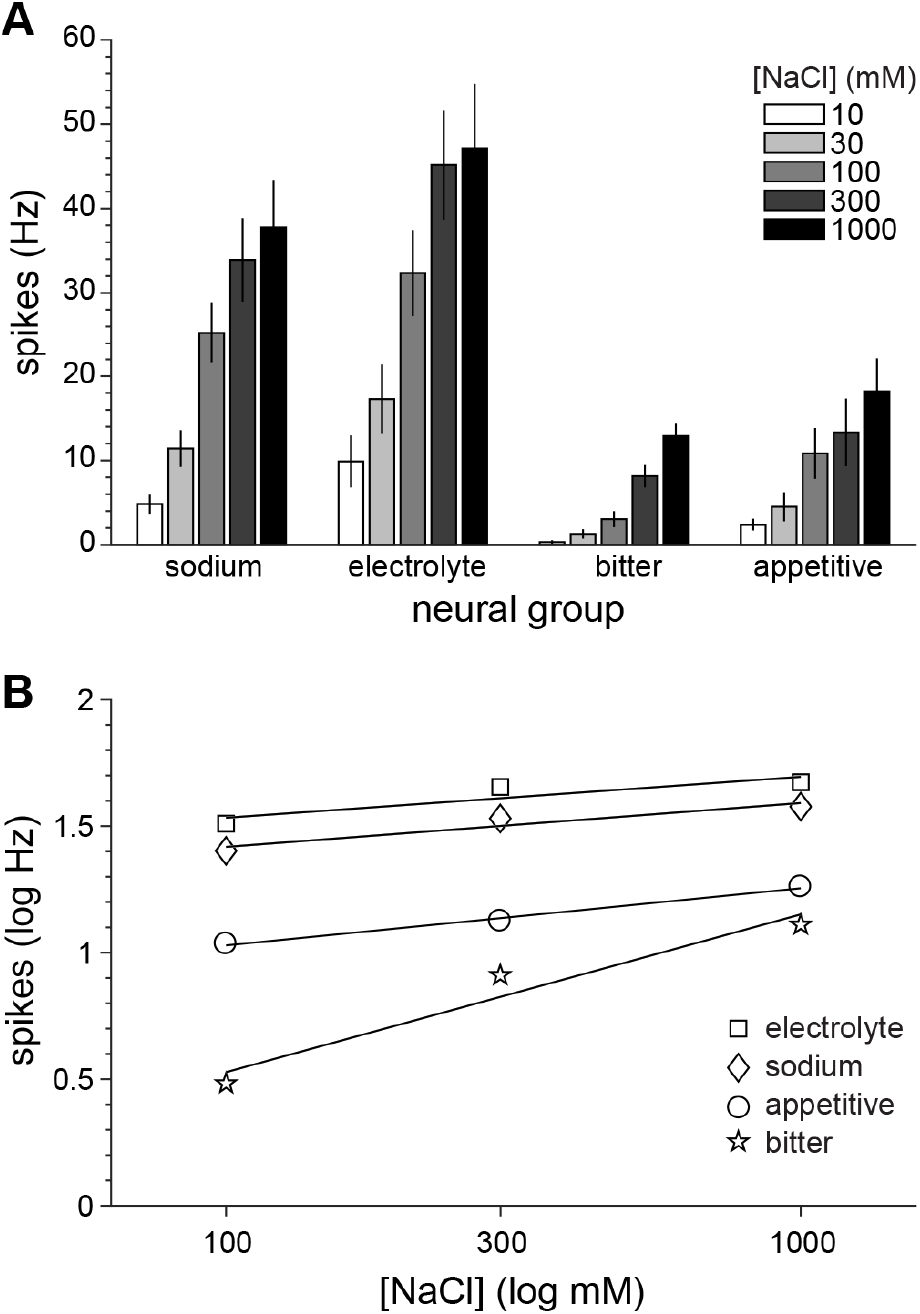
Activity to NaCI in neural groups. **A**, Mean ± s.e.m. firing in Hz by sodium (*n* = 12), electrolyte (*n* = 7), bitter (*n* = 15), and appetitive (*n* = 11) PbN neurons to a concentration series of NaCI (legend). **B**, Doubly logarithmic (base 10) plot of mean firing in Hz by each neural group (legend) to the three highest concentrations of NaCI from the series. Lines represent least-squares fits of mean values.

**Table 1.** Numbers of neurons of each type that significantly fired to each [NaCI]. For each class, the *P*-level is from a *χ*^2^ test (*df* = 4) of the null hypothesis that equal numbers of cells fired across concentrations. Parenthetical terms denote % of total cells for that type. *, significant at α = 0.05.

| Neural class | 10 mM | 30 mM | 100 mM | 300 mM | 1000 mM | $P$-level |
| --- | --- | --- | --- | --- | --- | --- |
| sodium | 9 (75%) | 10 (83%) | 12 (100%) | 12 (100%) | 12 (100%) | 0.95 |
| electrolyte | 5 (71%) | 6 (86%) | 7 (100%) | 7 (100%) | 7(100%) | 0.97 |
| bitter | 3 (20%) | 3 (20%) | 8 (53%) | 13 (87%) | 15 (100%) | 0.005* |
| appetitive | 4 (36%) | 6 (54%) | 10 (91%) | 10 (91%) | 11 (100%) | 0.34 |

That bitter neurons showed broad responsivity to unconditionally avoided bitter taste and salt chemicals suggests bitter cells are involved with aversive coding and agrees with the proposed co-activation of bitter sensing receptor cells by concentrated salts (Oka et al., 2013).

### Stimulation of the Vc activates PbN gustatory neurons

Taste-active PbN neurons in each group included cells that produced variable-latency spikes in response to weak electrical pulse stimulation of the orosensory Vc (V+ neurons; Figure 3A, 3B) and neurons that did not show reliable evoked activity to orosensory Vc stimulation (V− cells; Figure 3A, 3C). Latencies to evoked spikes in V+ PbN neurons varied with gustatory neuron class (Kruskal-Wallis *H*_3_ = 16.8, *P* = 0.001), with V+ bitter cells showing the longest median latency (Figure 3D) and a latency distribution that significantly differed from that of V+ sodium units (Dunn-Bonferroni adjusted pairwise comparison, *P* < 0.001). This trend may reflect that Vc-parabrachial projection neurons with different morphological or physiological properties transmit to different types of PbN gustatory cells. Considering total numbers (Table 2), the ratio of V+ to V− taste-active PbN neurons (2.3:1) was significantly larger than expected by chance (test of the null ratio of 1:1, *χ*^2^ = 10.2, *df* = 1, *P* = 0.001). Although the sampling of neurons was pseudorandom considering targeted placement of electrodes by coordinates, these data may suggest that areas of the PbN contain more taste-active neurons receiving contact from the Vc than not.

**Figure 3.**
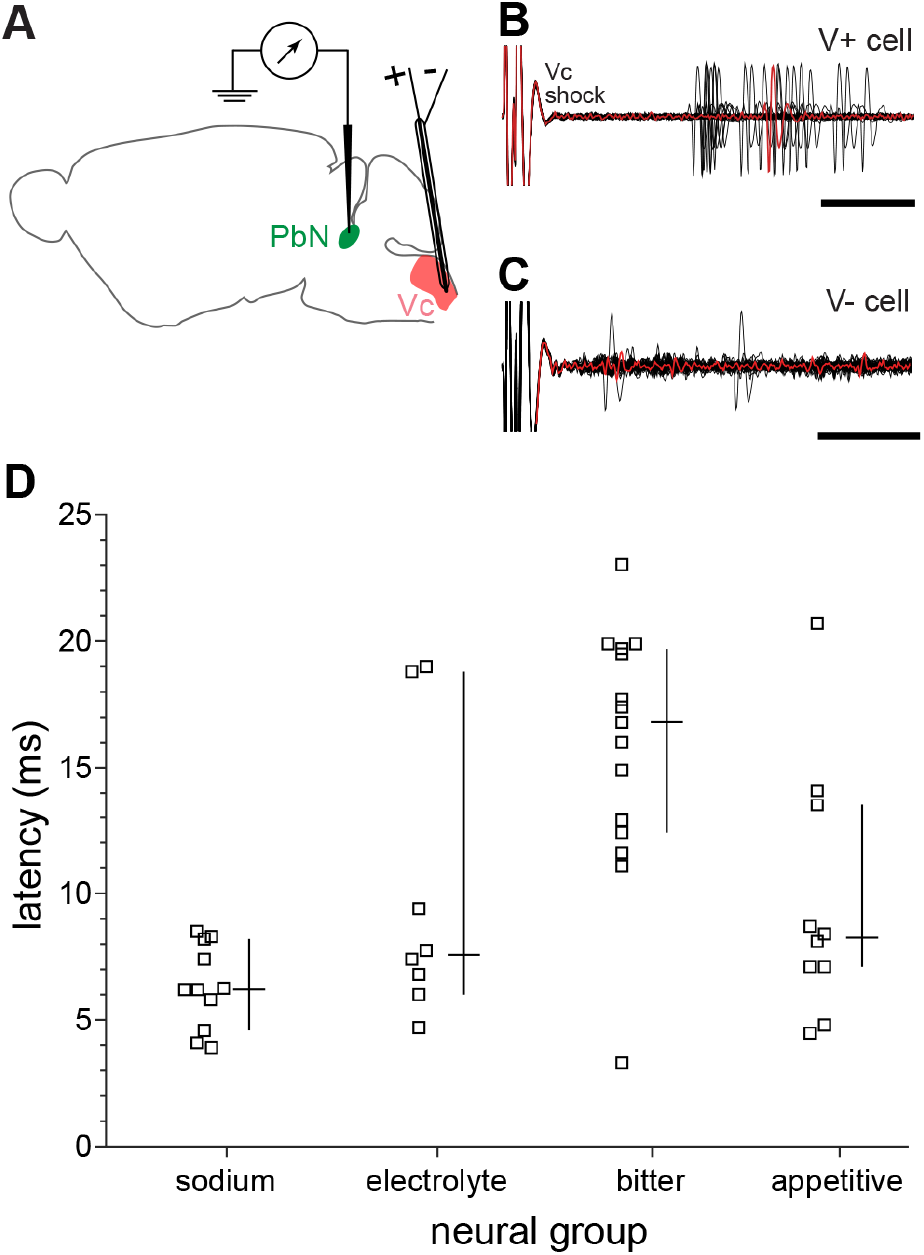
Investigation of electrophysiologic coupling between the Vc and taste-active PbN neurons. **A**, Schematic of experimental setup. Weak electrical pulses were delivered to an oral sensory region of the Vc while electrophys¡ologically monitoring for evoked spikes in PbN gustatory units. **B**, Raw electrophysiological sweeps showing an example PbN gustatory neuron that reliably spiked in response to electrical pulse stimulation of the Vc (V+ cell). C, Raw electrophysiological sweeps from an example PbN gustatory neuron that did not reliably spike to Vc stimulation (V− cell). For panels B and C, 40 sweeps aligned by the electrical stimulus (Vc shock) are superimposed, with an example single sweep shown in red. Scale bars = 5 ms. **D**, Latencies to fire to Vc stimulation for sodium (*n* = 11), electrolyte (*n* = 8), bitter (n = 15), and appetitive (*n* = 10) PbN neurons identified as V+. Plot column for each group shows latencies for individual neurons (left, squares), the median latency (right, horizontal line of cross), and the 95% Cl* of the median (right, vertical line of cross).

**Table 2.** Number of V+ and V− taste-active PbN neurons in each gustatory tuning class.

|  | sodium | electrolyte | appetitive | bitter | total |
| --- | --- | --- | --- | --- | --- |
| V+ | 11 | 9 | 10 | 16 | 46 |
| V- | 4 | 3 | 7 | 6 | 20 |

### Sensitivity to nociceptive stimuli in PbN gustatory neurons

Forty-eight taste-active PbN neurons successfully completed testing with all gustatory, thermal, and chemesthetic stimulus trials. From this sample, 33 (69%) were V+, with a subgroup of 9 V+ PbN gustatory cells identified to show notably strong, significant firing to the TRPV1 agonist CAP applied to the rostral-ipsilateral tongue. Neuronal responses to CAP were long-lasting, continuing after cessation of stimulus delivery and during oral application of a post-stimulus water rinse (e.g., Figure 4). Lingering influence is a hallmark of firing to lingual capsaicin by orosensory chemonociceptive Vc neurons (Carstens et al., 1998) and would be expected to arise in PbN cells receiving trigeminal-mediated CAP input.

**Figure 4.**
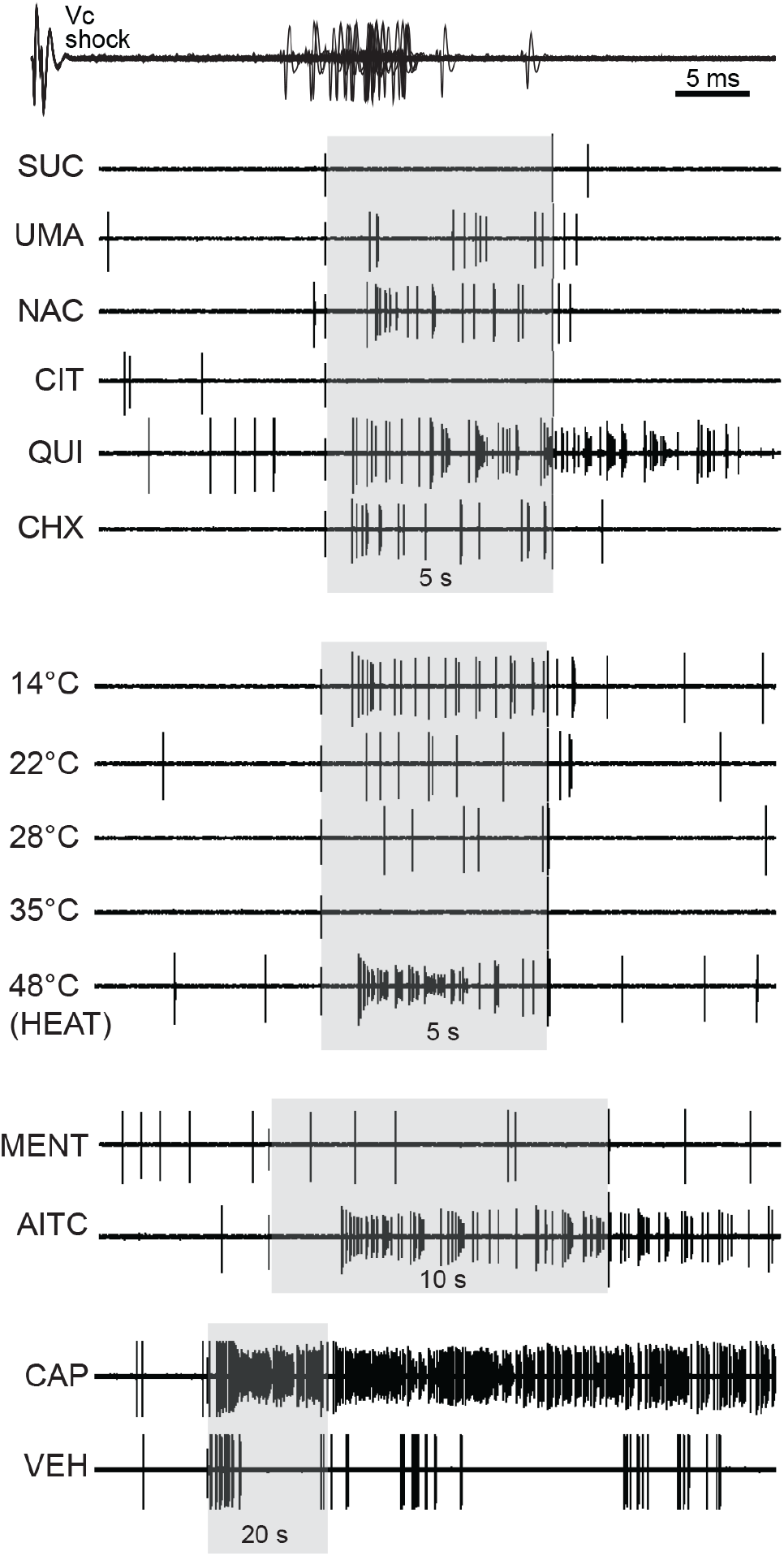
Raw electrophysiological sweeps showing activity by a single PbN neuron to electrical stimulation of the Vc (Vc shock, 40 sweeps superimposed) and oral delivery of taste stimuli (SUC, UMA, NAC, CIT, QUI, and CHX, abbreviated as in Figure 1), thermal-adjusted water (14°, 22°, 28°, 35°, and 48°C [HEAT], all measured as oral temperature), and chemesthetic stimuli (1.28 mM L-menthol [MENT], 1 mM allyl isothiocyanate [mustard oil, abbreviated AITC], 1 mM capsaicin [CAP], and the CAP vehicle [VEH]). Grayed areas denote oral stimulus application periods, denoted in s.

Six of the 9 CAP-responsive PbN gustatory neurons also displayed significant firing to oral delivery of the TRPA1 agonist AITC. This stimulus induced significant firing in only 3 additional CAP-unresponsive units out of the sample of 48 defined above. Thus, AITC predominantly excited CAP-responsive taste cells. Co-sensitivity to AITC and CAP in individual PbN neurons follows the frequent coexpression of TRPA1 and TRPV1 by somatosensory and trigeminal fibers (Kobayashi et al., 2005; Bautista et al., 2006) and partial activation of TRPV1 by millimolar concentrations of AITC (Everaerts et al., 2011). Further, AITC and also CAP induced, on average, a unique slow temporal rise in firing in CAP-responsive PbN gustatory neurons that thermal or taste stimuli (Figure 5). This observation agrees with the slow-building responses that lingual mustard oil and capsaicin induce in Vc neurons (Carstens et al., 1998) upstream of the PbN and supports the notion that responses to CAP and AITC in PbN taste neurons were of trigeminal origin. CAP-responsive gustatory cells showed only relatively poor or no sensitivity to MENT (Figure 5), which is a known stimulant of the TRPM8 ion channel expressed by a subset of trigeminal afferents (McKemy et al., 2002).

**Figure 5.**
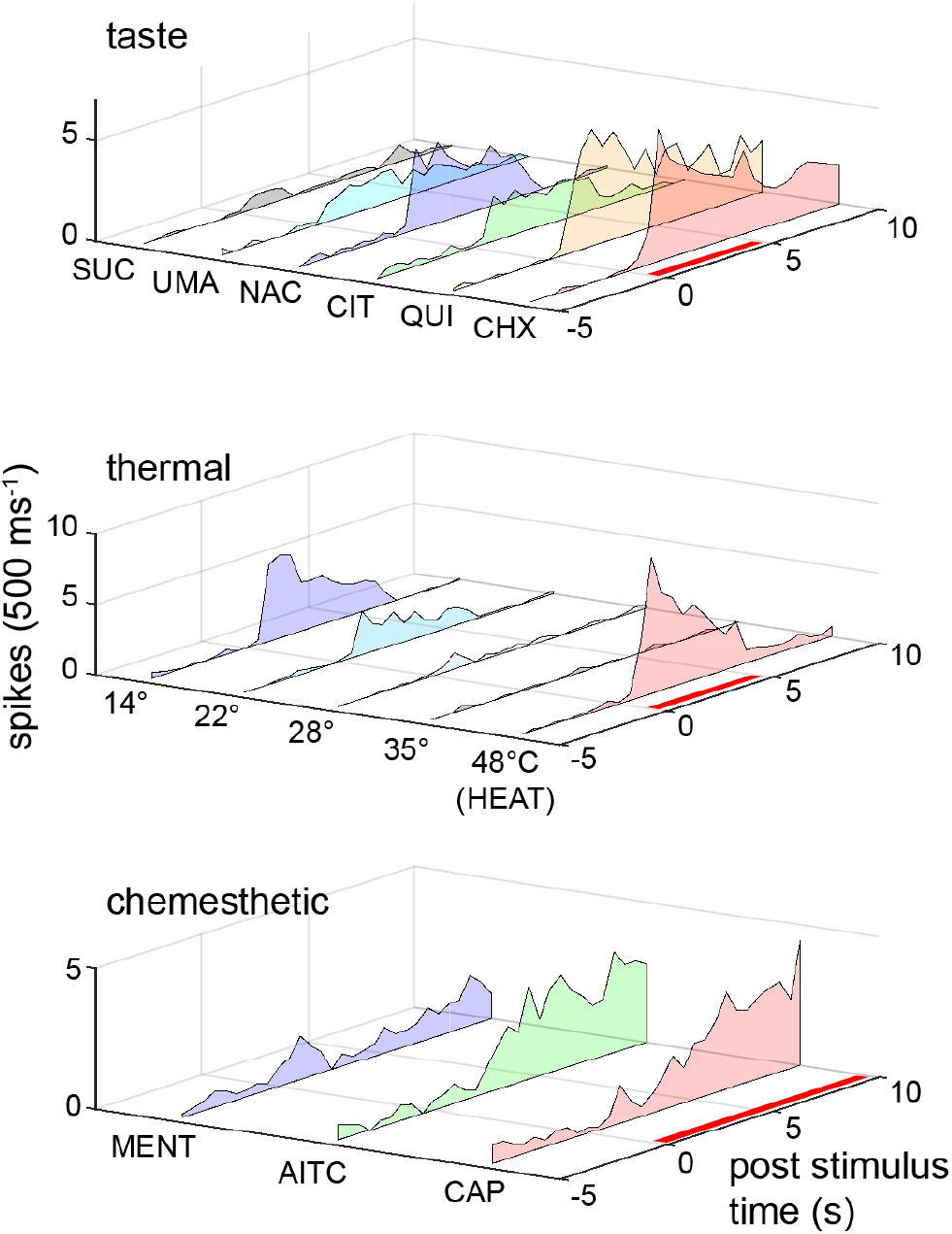
Peristimulus time histograms showing the mean time course of responding (in spikes per 500 ms) to taste, thermal, and chemesthetic stimuli (abbreviated as in Figures 1 and 4) across PbN gustatory neurons that significantly fired to oral delivery of capsaicin (*n* = 9). Red bar along the time axis denotes stimulus application period.

Finally, all 9 CAP-responsive PbN gustatory neurons significantly fired to HEAT. However, HEAT broadly activated the cellular population, significantly exciting 12 additional CAP-unresponsive neurons in the sample 48 cells tested with all taste, thermal, and chemesthetic trials. Although noxious heat engages the capsaicin receptor TRPV1 (Caterina et al., 1997), some decorrelation between sensitivity to HEAT and CAP is expected given evidence for involvement of non-TRPV1 mechanisms in heat detection. For instance, imaging data from mice on trigeminal ganglion neurons that fire to noxious heat demonstrated that nearly 30% are insensitive to capsaicin and that pharmacological blockade of TRPV1 reduces firing to high temperatures in only a minority of these cells (Yarmolinsky et al., 2016).

### An association between nociceptive and bitter activity in taste-active PbN neurons

Plotting the relationship between firing to nociceptive stimuli and cellular gustatory profiles revealed that the TRPV1 and TRPA1 agonists CAP and AITC primarily excited bitter-class PbN neurons – cells with unique high responsiveness to the bitters QUI and CHX (Figure 6) and elevated sensitivity to aversive concentrations of NaCI (Figure 2). Moreover, responses to CAP predominantly emerged in bitter taste neurons identified at V+ and did not arise in any V− cells (Figure 7A, 7B), implying upstream trigeminal coupling in PbN bitter taste units leads to their co-sensitivity to nociceptive stimuli. The same trend was noted for AITC, with the exception of a single V− cell that showed activity to this input (Figure 7B). HEAT was capable of more broadly stimulating V+, and some V−, PbN gustatory neurons of variable gustatory tuning, including bitter, electrolyte, and sodium units, albeit produced its largest responses, by far, in V+ bitter taste cells (Figure 7A, 7B). Cooling activated some V+ bitter cells, but more frequently excited electrolyte-responsive units (Figure 7A). Overall, these data implied that for V+ PbN taste neurons, excitatory firing to nociceptive heat and chemesthetic stimuli was positively associated with cellular co-sensitivity to bitter tastants.

**Figure 6.**
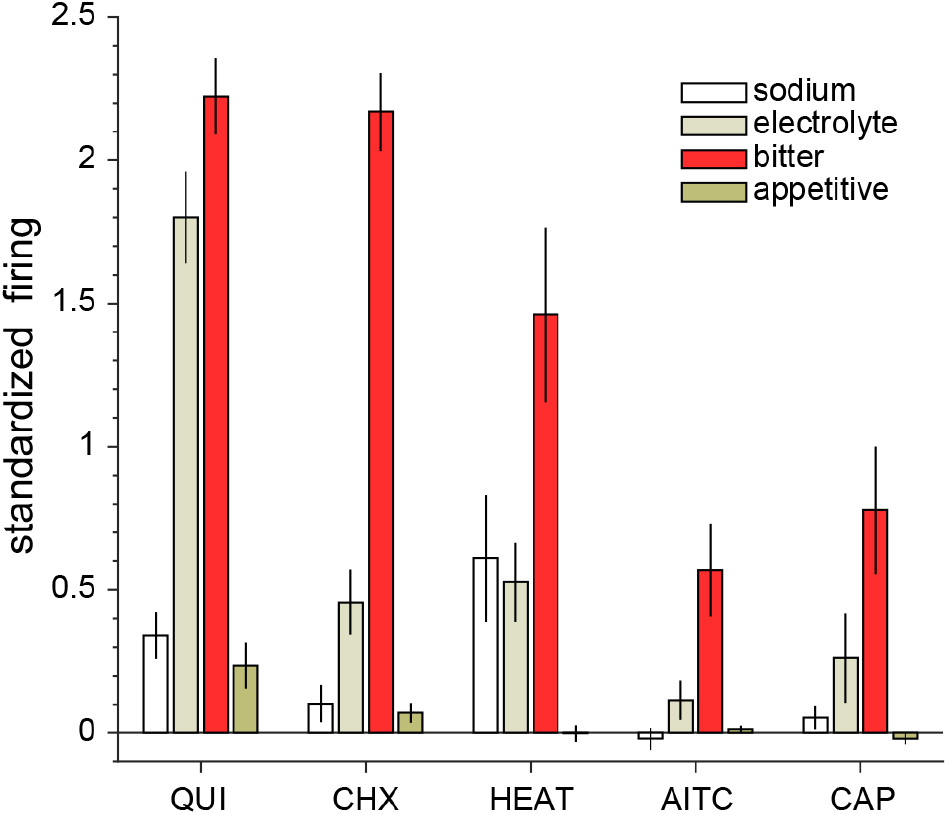
Mean ± s.e.m. standardized firing to oral delivery of the bitter tastants 10 mM quinine (QUI) and 0.1 mM cycloheximide (CHX) and the nociceptive stimuli 48°C (HEAT), 1 mM allyl isothiocyanate (mustard oil, abbreviated AITC), and 1 mM capsaicin (CAP) for sodium (*n* = 13), electrolyte (*n* = 9), bitter (*n* = 15), and appetitive (*n* = 12) PbN neurons. Note that firing to capsaicin is corrected for activity to the capsaicin vehicle.

**Figure 7.**
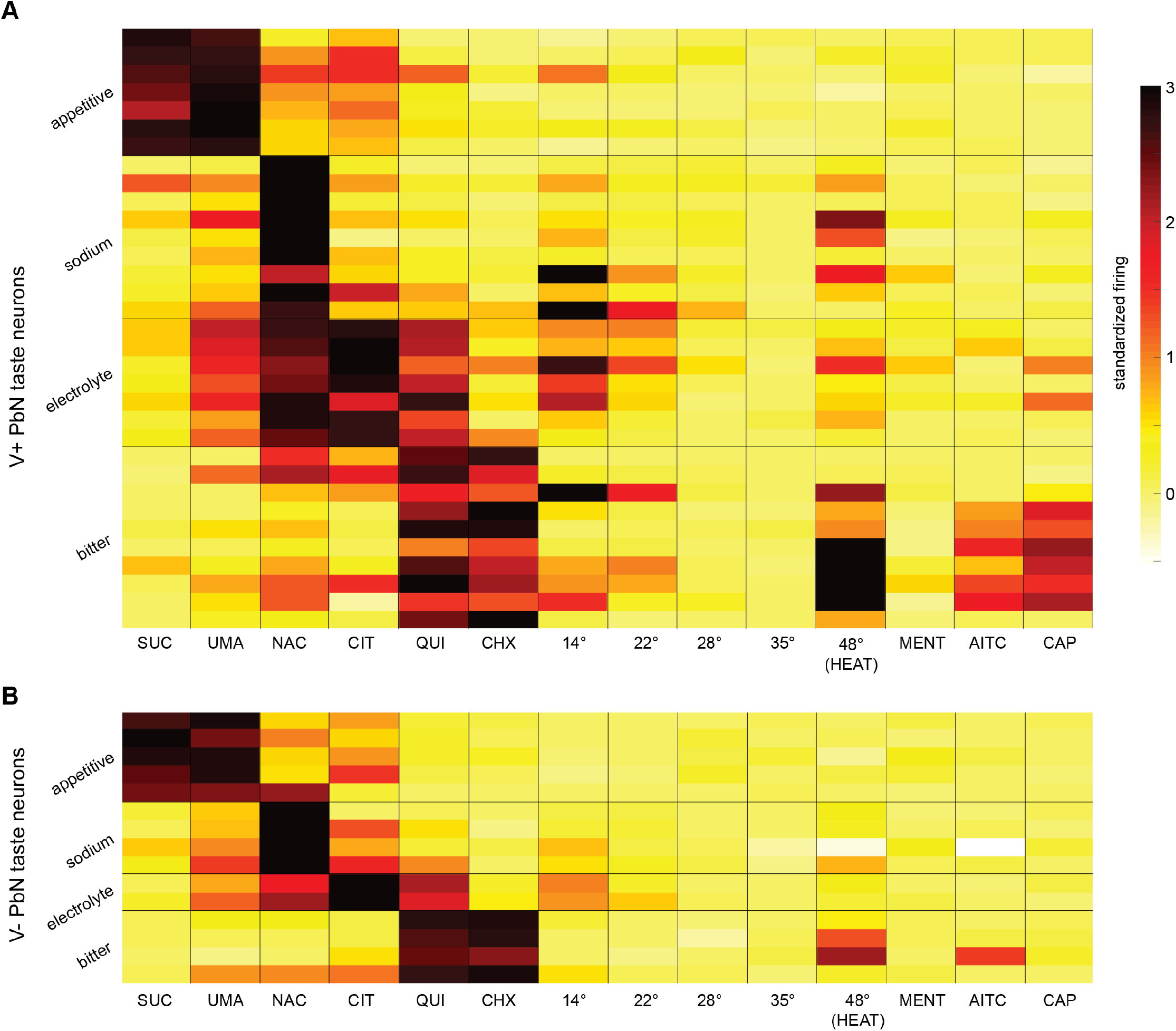
Heatmaps showing standardized firing to taste, thermal, and chemesthetic stimuli (abbreviated as in Figures 1 and 4) for individual appetitive, sodium, electrolyte, and bitter PbN neurons (each row gives response data for an individual neuron) that showed evidence of electrophysiological coupling with the Vc (**A**, V+ PbN taste neurons) or did not show such coupling (**B**, V− PbN taste neurons). Heat scale in A applies to both panels. Note that firing to capsaicin is corrected for activity to the capsaicin vehicle.

Closer inspection of activity across individual V+ PbN taste neurons suggested the correlation between nociceptive and bitter responses was partly bitter stimulus-dependent. For instance, although CAP activity was predominantly carried by bitter-class cells firing to QUI and CHX, CHX was a more selective stimulus for this cell type (Figure 7A). In comparison, QUI effectively stimulated bitter and also electrolyte neurons, with most cells in the latter group showing insensitivity to CAP (Figure 7A). Thus, although related to bitter signaling, firing to CAP appears to differentially associate with responding to the bitters QUI and CHX, showing a stronger tie to CHX.

This trend was quantitatively borne out in a multiple regression analysis. Variance in responding to CAP across taste-active V+ PbN neurons (*n* = 33) was correlated with activity to QUI and CHX (multiple *R* = 0.64, F_2,30_ = 10.1, *P* < 0.001). However, only firing to CHX showed a significant unique correlation with CAP activity (*sr* between firing to CAP and CHX [*s*_CAP×CHX_] = +0.48, *P* = 0.002; *sr*_CAP×QUI_ = +0.01, *P* = 0.9), which follows the unique association between responses to CAP and CHX visually apparent in plots of V+ cellular activity (Figure 7A). The same pattern was identified for activity to AITC, where firing to this input by V+ cells was associated with activity to both QUI and CHX (multiple *R* = 0.56, F2,30 = 6.8, *P* = 0.004) with only CHX contributing significantly to this relationship (*sr*_AITC×CHX_ = +0.41, *P* = 0.01; *sr*_AITC×QUI_ = +0.06, *P* = 0.7; see Figure 7A). Thus, sensitivity to the bitter tastant CHX in V+ PbN gustatory cells is a notable predictor of their responsiveness to chemesthetic stimuli associated with activation of the nocisensors TRPV1 and TRPA1.

Regression analyses also identified that HEAT activity in V+ PbN taste neurons was associated with activity to CHX (*R* = 0.39, *F*_1,31_ = 5.6, *P* = 0.03) but not both CHX and QUI, with the latter condition leading to a non-significant multiple *R*(*P* = 0.074). These effects agree with the broader ability of QUI and HEAT to stimulate multiple cellular types in addition to bitter-class neurons (Figure 7A).

Multiple regression analyses were not carried out for V− taste-active PbN neurons due to low sample size and the observation that somatosensory nociceptive and thermal firing was markedly blunted in V− compared to V+ taste-active cells (Figure 7). Altogether, the above data suggest there is a predictive relationship between activation to nociceptive inputs and bitter tastants in gustatory-active PbN cells that is tied to their receipt of input from upstream trigeminal nuclei.

### Trigeminal input contributes to nociceptive activity in PbN bitter taste neurons

To delineate whether Vc input was involved with sensitivity to nociceptive stimuli in trigeminal-coupled gustatory cells, responses to HEAT were recorded from V+ PbN bitter taste neurons during optogenetic-assisted photoinhibition of the orosensory Vc. This was accomplished using VGAT-ChR2 optogenetic mice, which afford laser-induced rapid and reversible suppression of oral sensory firing in the Vc (Figure 8A) while recording from PbN neurons. Overall, 146 responses to brief applications of HEAT were acquired from 8 PbN cells under laser-on (Vc laser) and laser-off (control) conditions. Raw data from application of this procedure to a single PbN unit are shown in Figure 8B, where spike firing to HEAT was found to be visibly suppressed over 10 Vc laser compared to 10 control trials. For this unit, responses in Hz to HEAT were always lower on Vc laser trials, resulting in an auROC of 1.

**Figure 8.**
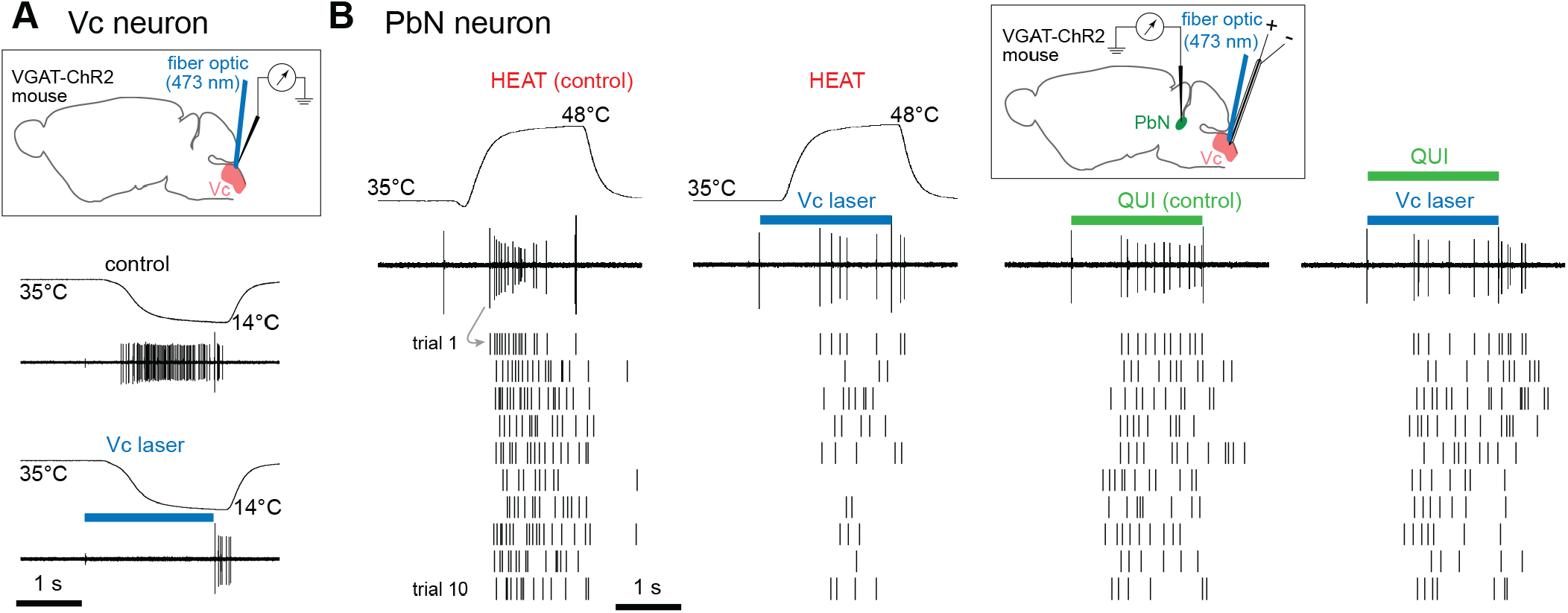
Example effects of optogenetic-assisted suppression of oral sensory neural activity in VGAT-ChR2 mice. **A**, Raw electrophysiological sweeps showing firing by a Vc neuron in a VGAT-ChR2 mouse to an oral cooling ramp (shown, measured as oral temperature) presented by itself (control trial) or in unison with blue laser light delivery to the Vc (Vc laser trial), which engaged local inhibitory neurons. Spiking by this cell to cooling was notably strong under the control condition but suppressed on the Vc laser trial; note spiking recovered immediately following removal of light (bottom trace). Inset shows experiment schematic, where the recording electrode and fiber optic for laser delivery both targeted the same area of the Vc. Note that Vc unit recording was not part of the present analyses but is shown here to illustrate the optogenetic inhibitory effect. **B**, Activity by a V+ PbN bitter taste neuron in a VGAT-ChR2 mouse during oral presentations of noxious heat (48°C, HEAT) over 10 control and 10 Vc laser trials (two leftmost columns), and 10 mM quinine (QUI) over 10 control and 10 Vc laser trials (two rightmost columns). The raw electrophysiological sweep for the first trial of each stimulus/condition is shown to illustrate conversion of neurophysiological data to raster spikes; thermal traces on HEAT trials reflect oral temperature. For this PbN neuron, firing to HEAT was markedly and repeatedly suppressed on Vc laser compared to control trials, whereas quinine activity did not differ between conditions; see results for details. Inset shows experiment schematic, where the recording electrode was positioned in the PbN while the fiber optic for light delivery and the electrical stimulation electrode both targeted the same oral sensory-verified region of the Vc.

ROC permutation tests applied to 7 individual units revealed 5 showed significantly reduced firing to HEAT over 10 Vc laser compared to 10 control trials (*P* < 0.05; Figure 9A). Permutation statistics could not be computed for the one remaining neuron due to acquisition of only 3 trials per laser-on and control conditions. However, HEAT activity by this cell was always lower on Vc laser trials (auROC = 1). Thus, in VGAT-ChR2 mice, Vc photoinhibition suppressed firing to HEAT in 6 out of 8 (75%) V+ taste-active PbN cells tested. Moreover, several of these neurons showed reduced median firing to HEAT on Vc laser trials compared to control, with non-overlapping 95% Cl*s between conditions (Figure 9B) implying such reductions were significant. While activity to HEAT did show reduction, it was not completely suppressed (i.e., to zero spikes) by Vc photoinhibition in most cases, potentially reflecting network complexity.

**Figure 9.**
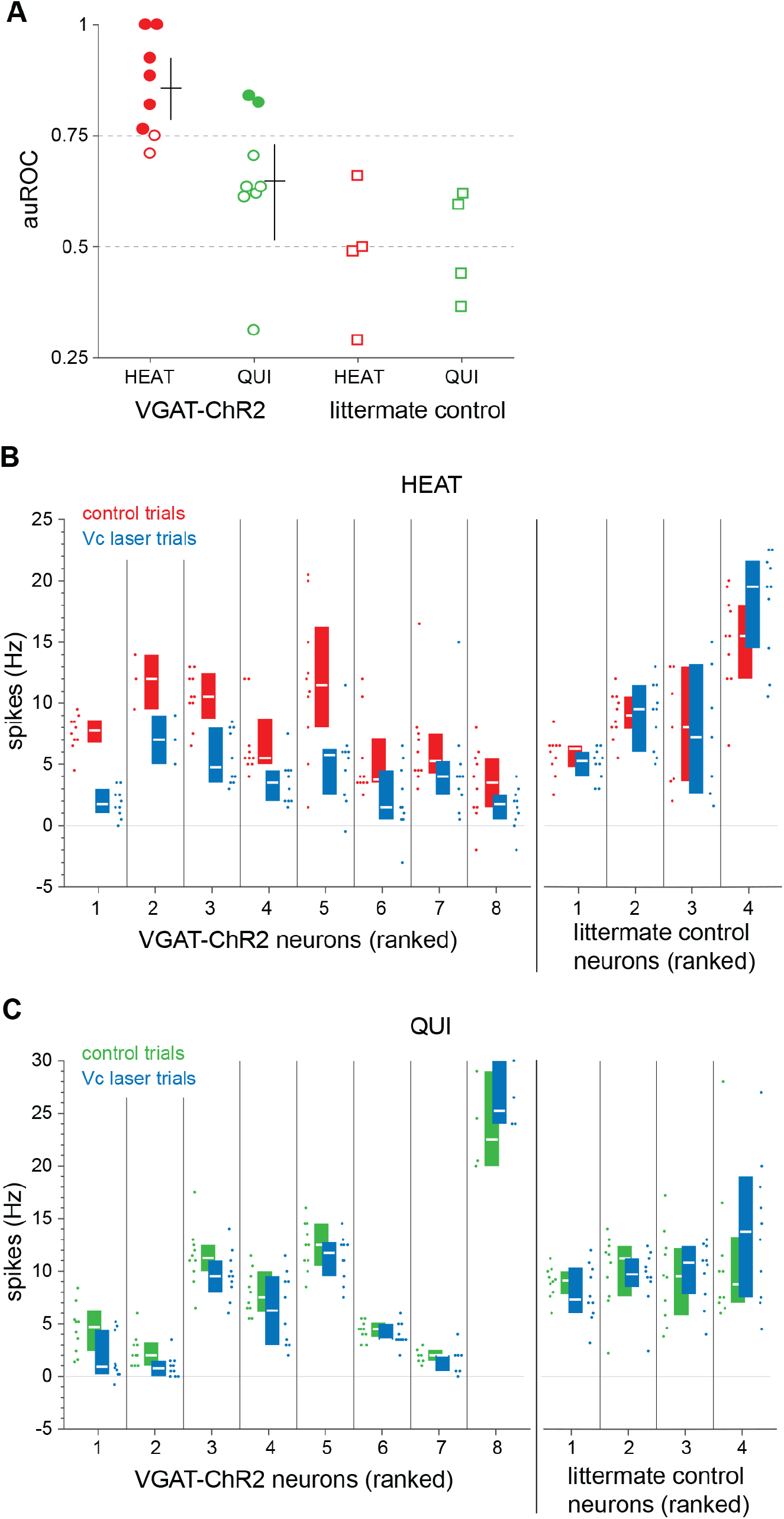
Analyses of firing to noxious heat (48°C, HEAT) and 10 mM quinine (QUI) for multiple Vc-coupled PbN bitter neurons during optogenetic-assisted inhibition targeted to the Vc. **A**, Plotted are auROC values for individual PbN neurons from VGAT-ChR2 mice (circles) and non-optogenetic littermate mice (squares) computed from their firing to HEAT (red) or QUI (green) under control and Vc laser conditions. Filled markers represent auROC values that equal 1 or significantly differ from chance; open markers indicate;? auROC values that do not differ from chance. For neurons acquired from VGAT-ChR2 mice, the median auROC value (horizontal line of cross) and 95% Cl* of the median (vertical line of cross) is ? shown to the right of their auROC scores for HEAT and QUI. **B**, Each numbered column shows for an individual PbN neuron all responses in Hz to HEAT (points) and the 95% Cl* of the median HEAT response (vertical bar spans the interval, with the median marked by a white horizontal line) on control (red) and Vc laser (blue) trials. Neurons from VGAT-ChR2 and non-optogenetic littermate mice are ranked along the abscissa in descending order of their respective auROC values for HEAT, shown in panel A. C, Numbered columns show for individual neurons all responses in Hz to QUI (points) and the 95% Cl* of the median QUI response (as in panel B) on control (green) and Vc laser (blue) trials. Neurons are ranked as in panel B but based on auROC values for QUI.

To evaluate the stimulus specificity of the optogenetic effect, responses to QUI were recorded during Vc laser and control trials from 6 of the above PbN neurons and also 2 additional V+ PbN bitter taste units (*n* = 8 total) in VGAT-ChR2 mice. Six cells completed 10 pairs of these trials; 2 cells completed 4 and 7 trial pairs, respectively, making 142 QUI trials in total available for analysis. In contrast to its reduced firing to HEAT on Vc laser trials, the example PbN neuron in Figure 8B gave visibly similar responses to QUI over 10 Vc laser and 10 control trials, with ROC permutation statistics revealing no significant difference in firing to QUI between conditions (auROC = 0.62, *P* > 0.05). Overall, ROC permutation tests identified firing to QUI did not differ between Vc laser and control trials for 6 out of 8 neurons (75%) recorded from VGAT-ChR2 mice (*P* > 0.05; Figure 9A). Effects in the two remaining cells appeared minor given their low firing rates to QUI on several control trials (VGAT-ChR2 neurons 1 and 2 in Figure 9C). Moreover, all VGAT-ChR2 neurons showed 95% Cl*s for median firing to QUI that overlapped between Vc laser and control conditions (Figure 9C), which differed from the non-overlap of these intervals observed for VGAT-ChR2 cellular activity to HEAT (Figure 9B).

Finally, to control for non-specific laser effects on brain tissue, responses to HEAT and QUI were recorded from bitter-class V+ PbN taste neurons (*n* = 4) in littermate optogenetic-negative mice during Vc laser and control trials. All PbN neurons sampled from optogenetic-negative mice completed 10 pairs of these trials for QUI (80 QUI trials total); for HEAT, 3 PbN neurons completed 10 Vc laser on/off trial pairs, whereas 1 cell completed 7 pairs (74 HEAT trials overall). ROC permutation tests applied to individual neurons revealed no differences in firing to HEAT or QUI between laser-on and control conditions in non-optogenetic animals (*P* > 0.05; Figure 9A). Moreover, all cells acquired from these mice showed 95% Cl*s for median activity to HEAT and QUI that overlapped between Vc laser and control trials (Figure 9B, 9C), unlike VGAT-ChR2 cellular responses to HEAT (Figure 9B). These results attributed the light-induced suppression of PbN sensory firing observed in VGAT-ChR2 mice to optogenetic activation of Vc inhibitory circuits.

Altogether, the above data show temporary suppression of neural activity in an orosensory region of the Vc differentially impacts PbN unit responses to nociceptive and bitter taste stimulation. Specifically, photoinhibition of Vc circuits caused a reliable and predominant reduction in firing to HEAT compared to QUI in a subset of Vc-coupled PbN bitter taste neurons. This finding implies trigeminal pathways contribute to nociceptive activity in taste-active PbN cells.

### Nociceptive and aversive taste signals converge in the lateral PbN

Histological reconstruction of recording sites for 40 taste-active PbN neurons identified that the numbers of V+ (n = 15) and V− (*n* = 12) cells acquired from the dorsal lateral, central lateral, and medial PbN did not reliably differ (*χ*^2^ = 0.33, *df* = 1, *P* = 0.56, Figure 10). Cells of all gustatory types were found in these regions.

**Figure 10.**
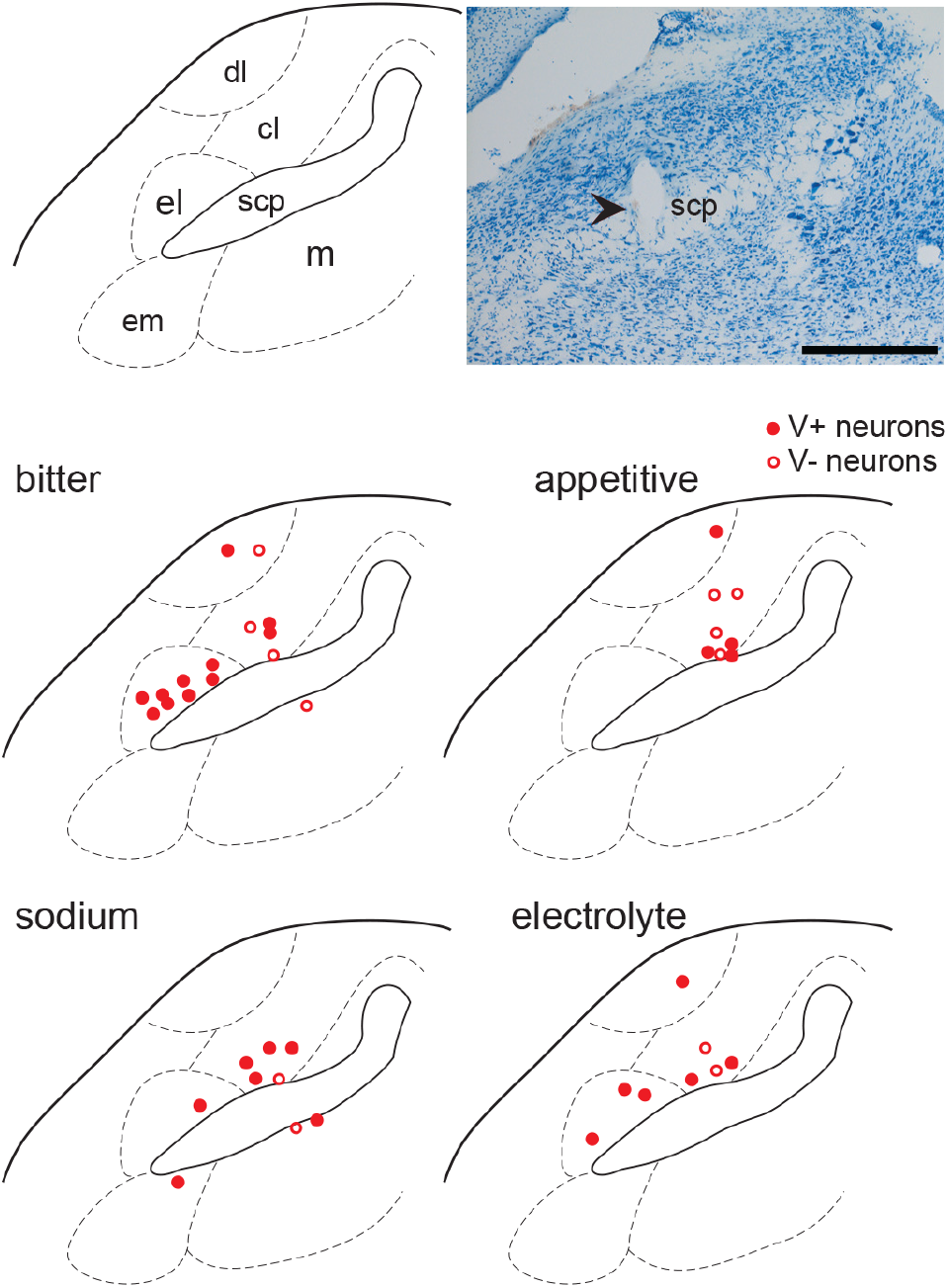
Histological analysis of recording sites (red circles) for bitter (*n* = 15), appetitive (*n* = 8), sodium (*n* = 9), and electrolyte (*n* = 8) PbN neurons. Legend denotes whether recorded cells were identified as V+ or V−. Photograph depicts electrolytic lesion (arrowhead) made in the lateral PbN where a bitter-class neuron was acquired. Scalebar = 500 μm. Reconstructed lesion sites are plotted on only one coronal plane of the PbN for simplicity; actual lesion sites were observed approximately 60 μm around this rostro-caudal location. Abbreviations: dorsal lateral nucleus (dl); central lateral nucleus (cl); external lateral PbN (el); external medial PbN (em); medial parabrachial area (m); superior cerebellar peduncle (scp).

Yet of the 12 neurons observed in the external lateral PbN, all were V+, with 8 of these cells identified as bitter-class neurons that co-fired to nociceptive stimuli, including CAP and HEAT. With one exception, the remaining sodium and electrolyte V+ taste-active cells in the external lateral PbN responded to oral presence of HEAT. Thus, lateral parabrachial circuits contain neurons sensitive to avoided taste and oral nociceptive inputs.

## Discussion

Electrical stimulation of the orosensory Vc excited multiple types of parabrachial gustatory neurons. We show here that a subpopulation of these cells responsive to the bitter taste stimuli quinine and cycloheximide, and with elevated sensitivity to high (aversive) salt, co-fired to oral delivery of agonists of the nocisensors TRPV1 and TRPA1 on somatosensory fibers, including capsaicin, mustard oil, and noxious heat. Firing to cycloheximide was an important identifier of such neurons, given broader activation of bitter and other cells by quinine. Further, acute photoinhibition of Vc circuits reversibly suppressed nociceptive heat activity in a subset of bitter-active PbN cells. Thus, trigeminal pathways appear to contribute to oral nociceptive firing in PbN bitter taste neurons. Such neurons would be capable of serving roles in protective coding beyond only taste.

Although our results imply trigeminal sensory signals reach parabrachial gustatory units, it is important to consider potential alternative mechanisms. Earlier data from rats suggest millimolar (i.e., taste-like) concentrations of quinine can electrophysiologically excite and inhibit trigeminal lingual nerve fibers in vivo (Pittman and Contreras, 1998) and increase intracellular calcium ([Ca^2+^]i) in cultured capsaicin-sensitive trigeminal ganglion neurons at room temperature (Liu and Simon, 1998). Such data would imply that presynaptic trigeminal neurons mediate both nociceptive and bitter activity in PbN cells co-responsive to these stimuli. Yet a different picture emerges from neuronal data on the Vc, which was presently linked to nociceptive firing in PbN bitter taste cells. Electrophysiological recordings in mice revealed oral somatosensory Vc neurons are unresponsive to temperature-controlled taste stimuli, including millimolar quinine, that activate gustatory cells in the NTS (Lemon et al., 2016). Further, Simons et al. (2003b) used c-Fos labeling of cellular activity and electrophysiology in rats to ask whether lingual presence of an elevated concentration of quinine could excite oral sensory Vc neurons, including chemonociceptive capsaicin-sensitive units, but found no response. Thus, Vc neurons are insensitive to quinine, implying Vc-coupled PbN bitter-class neurons receive quinine signals from the gustatory NTS, with somatosensory input contributing to their firing to nociceptive stimuli such as tongue-applied capsaicin. The converse that capsaicin could reliably excite peripheral gustatory afférents is not strongly supported (Dahl et al., 1997; Simons et al., 2003a; Boucher et al., 2014).

An additional mechanism for bitter-pain overlap is apparent in the afferent system of the solitary chemosensory cells (SCCs) of the airways. SCCs express some Tas2R bitter taste receptors and contact peptidergic fibers of nasal trigeminal and vagal nerves (Finger et al., 2003; Tizzano et al., 2011). While this arrangement could lead to co-firing to bitter and nociceptive stimuli in downstream neurons, involvement of this system with the present results is unclear. The expression of SCCs in the airways is outside of the intended scope of our study, albeit taste solutions delivered whole-mouth may have reached SCC-containing pharyngeal fields innervated by the superior laryngeal (SL) nerve (Tizzano et al., 2011). Nevertheless, SL fibers in mice respond to water and do not increase firing to quinine (Ohkuri et al., 2012). Cycloheximide was reported to raise [Ca^2+^]_i_ in rodent vomeronasal SCCs (Ogura et al., 2010) and excite nasal trigeminal fibers in vivo (Finger et al., 2003), but at concentrations 25 and 100 times greater, respectively, than the presently used 0.1 mM. In mice, 0.1 mM cycloheximide causes aversion in taste-salient behavioral assays (Boughter et al., 2005) and excites gustatory neurons in the NTS (Wilson et al., 2012). Other data show nasal SCCs do not express a taste receptor for cycloheximide (Finger et al., 2003) and do not increase [Ca^2+^]_i_ in response to up to 20 mM cycloheximide, which caused decreased [Ca^2+^]_i_ or cell death (Gulbransen et al., 2008).

Presently, Vc photoinhibition suppressed firing to noxious heat in an imperfect and variable manner across PbN bitter taste neurons. This may reflect an ability of heat to partly engage pathways untargeted by our optogenetic manipulation. As a further limitation to this approach, sudden perturbation of Vc input to the PbN may disrupt processing of off-target projections arriving in the parabrachial area (Otchy et al., 2015) including gustatory afferents, which may account for the minor reductions in quinine activity noted during Vc inhibition. Nevertheless, that inhibition focused to the Vc predominantly suppressed firing to noxious heat compared to quinine implies separate pathways are involved with transmitting these signals to common PbN neurons.

### The PbN and taste-somatosensory overlap

Electrical excitation of the Vc stimulated taste-active neurons in multiple parabrachial areas, but V+ neurons that fired to taste and nociceptive input, including a density of bitter-nociceptive cells, were found in the external region of the lateral PbN. Anatomical data show the lateral PbN receives afferent projections from the Vc (Cechetto et al., 1985; Bernard et al., 1989). Efferent projections of external and lateral PbN areas include the central nucleus of the amygdala, with parabrachioamygdaloid neurons firing to noxious thermal and mechanical stimuli delivered to trigeminal-supplied orofacial receptive fields (Bernard and Besson, 1990). The external and lateral PbN subnuclei also receive input from the NTS, including its caudal visceroceptive regions (Ricardo and Koh, 1978; Bernard et al., 1989) and areas mediating signals for aversive taste. In rats, external medial and external lateral PbN neurons show c-Fos expression following intraoral infusion of quinine (Yamamoto et al., 1994) and fire to application of bitters, including quinine and cycloheximide, to palate and posterior oral taste papillae innervated by, respectively, the facial and glossopharyngeal nerves, which supply gustatory input to NTS neurons presynaptic to the PbN (Halsell and Travers, 1997; Geran and Travers, 2009). Altogether, these data point to the PbN as a brain site where aversive taste and nociceptive processing would overlap, as discovered presently.

The present nociceptive-active bitter taste neurons in the lateral PbN would appear to convey a common aversive or protective signal serving multiple modalities as opposed to only taste. Involvement of lateral PbN cells with general protective processing is also supported by data showing this brain region links pain and avoidance circuits from different areas of the body to limbic structures. In rats, noxious heat and pinch applied to the hind paws and tail, supplied by spinal nerves, can activate lateral PbN parabrachioamygdaloid neurons co-sensitive to nociceptive stimulation of orofacial trigeminal fields (Bernard and Besson, 1990). Furthermore, lateral PbN neurons that express calcitonin gene-related peptide (CGRP), implicated to contribute to the parabrachioamygdaloid circuit (Schwaber et al., 1988), can respond to malaise, itch, extended food consumption, and unconditioned and conditioned aversive stimuli (Campos et al., 2018). Recent papers discuss lateral PbN CGRP cells as a neural mechanism encoding a general “stop” signal to ward off harm and posit this cellular population would also fire to aversive taste (Campos et al., 2018; Palmiter, 2018). Although the present work did not index the genetic signature of nociceptive-active bitter cells, their presence in the external lateral PbN and upstream coupling with the Vc suggest they may be part of the lateral PbN cellular population that transmits to the amygdala and is involved with general aversive coding.

Bitter taste activity in external lateral PbN cells may stem from projections from NTS regions that receive lingual-tonsillar glossopharyngeal nerve fibers. These fibers innervate caudal tongue gustatory receptors and terminate at the level of the NTS that meets the fourth ventricle, which contains cells projecting to the external lateral PbN (Hamilton and Norgren, 1984; Bernard et al., 1989; Herbert et al., 1990). Notably, glossopharyngeal fibers show robust firing to lingual presence of cycloheximide (Danilova and Hellekant, 2003), which was presently identified as a key predictor of nociceptive-active, bitter-class PbN neurons. That glossopharyngeal nerve input may drive bitter activity in these units further supports their involvement in protective coding, as severing the glossopharyngeal nerve impairs rejective, but not discriminative, rat oral behavioral responses to bitter stimuli (St. John and Spector, 1998; Geran and Travers, 2011).

Several questions remain concerning taste-somatosensory integration in the PbN. Vc pulse stimulation excited V+ taste-active PbN neurons sensitive to oral cooling. Yet the cooling agent menthol, an agonist of TRPM8 on somatosensory fibers, only poorly stimulated Vc-coupled PbN cooling units. Whether cooling signals in these cells are contributed by the subpopulation of menthol-insensitive, cooling-responsive trigeminal sensory neurons (Bautista et al., 2007) awaits direct investigation. Notably, some PbN neurons, particularly appetitive-class cells, did not fire to somatosensory stimuli but generated spikes to Vc pulse stimulation. The function of trigeminal coupling to these cells remains unclear, but, speculatively, may come online under conditions presently untested.

In closing, the present data reveal that neural messages associated with bitterness and oral pain are partly relayed to PbN neurons dually linked to gustatory and trigeminal processing. Our results agree with the postulate that PbN circuits mediate multi-system convergence associated with general protective coding and implicate classically-identified “gustatory” neurons in this process. Along with recent findings on involvement of taste neurons in crossmodal signaling (Fortis-Santiago et al., 2010; Escanilla et al., 2015; Maier et al., 2015), including multisensory valence coding (Samuelsen and Fontanini, 2017), our results highlight that understanding the circuit properties and broader sensory repertoire of gustatory-active brain cells is needed to delineate their roles in taste processing and beyond.

## Acknowledgements

The authors thank Brad Heldmann and Jordan Norris for assistance with histology. Supported by NIH grant DC 011579 to C.H.L.

## References

Abe J, Hosokawa H, Okazawa M, Kandachi M, Sawada Y, Yamanaka K, Matsumura K, Kobayashi S (2005) TRPM8 protein localization in trigeminal ganglion and taste papillae. Brain Res Mol Brain Res 136:91–98.

Bautista DM, Siemens J, Glazer JM, Tsuruda PR, Basbaum AI, Stucky CL, Jordt SE, Julius D (2007) The menthol receptor TRPM8 is the principal detector of environmental cold. Nature 448:204–208.

Bautista DM, Jordt SE, Nikai T, Tsuruda PR, Read AJ, Poblete J, Yamoah EN, Basbaum AI, Julius D (2006)TRPA1 mediates the inflammatory actions of environmental irritants and proalgesic agents. Cell 124:1269–1282.

Bernard JF, Besson JM (1990) The spino(trigemino)pontoamygdaloid pathway: electrophysiological evidence for an involvement in pain processes. J Neurophysiol 63:473–490.

Bernard JF, Peschanski M, Besson JM (1989) A possible spino (trigemino)-ponto-amygdaloid pathway for pain. Neurosci Lett 100:83–88.

Boucher Y, Simons CT, Carstens MI, Carstens E (2014) Effects of gustatory nerve transection and/or ovariectomy on oral capsaicin avoidance in rats. Pain 155:814–820.

Boucher Y, Simons CT, Faurion A, Azerad J, Carstens E (2003) Trigeminal modulation of gustatory neurons in the nucleus of the solitary tract. Brain Res 973:265–274.

Boughter JD, Jr., Raghow S, Nelson TM, Munger SD (2005) Inbred mouse strains C57BL/6J and DBA/2J vary in sensitivity to a subset of bitter stimuli. BMC Genet 6:36.

Braud A, Vandenbeuch A, Zerari-Mailly F, Boucher Y (2012) Dental afferents project onto gustatory neurons in the nucleus of the solitary tract. J Dent Res 91:215–220.

Campos CA, Bowen AJ, Roman CW, Palmiter RD (2018) Encoding of danger by parabrachial CGRP neurons. Nature 555:617–622.

Carstens E, Kuenzler N, Handwerker HO (1998) Activation of neurons in rat trigeminal subnucleus caudalis by different irritant chemicals applied to oral or ocular mucosa. J Neurophysiol 80:465–492.

Caterina MJ, Schumacher MA, Tominaga M, Rosen TA, Levine JD, Julius D (1997) The capsaicin receptor: a heat-activated ion channel in the pain pathway. Nature 389:816–824.

Caterina MJ, Leffler A, Malmberg AB, Martin WJ, Trafton J, Petersen-Zeitz KR, Koltzenburg M, Basbaum AI, Julius D (2000) Impaired nociception and pain sensation in mice lacking the capsaicin receptor. Science 288:306–313.

Cechetto DF, Standaert DG, Saper CB (1985) Spinal and trigeminal dorsal horn projections to the parabrachial nucleus in the rat. J Comp Neurol 240:153–160.

Chang FC, Scott TR (1984) Conditioned taste aversions modify neural responses in the rat nucleus tractus solitarius. J Neurosci 4:1850–1862.

Chase SM, Young ED (2007) First-spike latency information in single neurons increases when referenced to population onset. Proc Natl Acad Sci U S A 104:5175–5180.

Dahl M, Erickson RP, Simon SA (1997) Neural responses to bitter compounds in rats. Brain Res 756:22–34.

Danilova V, Hellekant G (2003) Comparison of the responses of the chorda tympani and glossopharyngeal nerves to taste stimuli in C57BL/6J mice. BMC Neurosci 4:5.

Dhaka A, Earley TJ, Watson J, Patapoutian A (2008) Visualizing cold spots: TRPM8-expressing sensory neurons and their projections. J Neurosci 28:566–575.

Ellingson JM, Silbaugh BC, Brasser SM (2009) Reduced oral ethanol avoidance in mice lacking transient receptor potential channel vanilloid receptor 1. Behav Genet 39:62–72.

Escanilla OD, Victor JD, Di Lorenzo PM (2015) Odor-taste convergence in the nucleus of the solitary tract of the awake freely licking rat. J Neurosci 35:6284–6297.

Everaerts W, Gees M, Alpizar YA, Farre R, Leten C, Apetrei A, Dewachter I, van Leuven F, Vennekens R, De Ridder D, Nilius B, Voets T, Talavera K (2011) The capsaicin receptor TRPV1 is a crucial mediator of the noxious effects of mustard oil. Curr Biol 21:316–321.

Eyiam S, Spector AC (2002) The effect of amiloride on operantly conditioned performance in an NaCl taste detection task and NaCl preference in C57BL/6J mice. Behav Neurosci 116:149–159.

Felizardo R, Boucher Y, Braud A, Carstens E, Dauvergne C, Zerari-Mailly F (2009) Trigeminal projections on gustatory neurons of the nucleus of the solitary tract: a double-label strategy using electrical stimulation of the chorda tympani and tracer injection in the lingual nerve. Brain Res 1288:60–68.

Finger TE, Bottger B, Hansen A, Anderson KT, Alimohammadi H, Silver WL (2003) Solitary chemoreceptor cells in the nasal cavity serve as sentinels of respiration. Proc Natl Acad Sci U S A 100:8981–8986.

Fortis-Santiago Y, Rodwin BA, Neseliler S, Piette CE, Katz DB (2010) State dependence of olfactory perception as a function of taste cortical inactivation. Nat Neurosci 13:158–159.

Franklin K, Paxinos G (2008) The mouse brain in stereotaxic coordinates, 3rd Edition. San Diego: Academic Press.

Gadziola MA, Wesson DW (2016) The Neural Representation of Goal-Directed Actions and Outcomes in the Ventral Striatum’s Olfactory Tubercle. J Neurosci 36:548–560.

Gauriau C, Bernard JF (2002) Pain pathways and parabrachial circuits in the rat. Exp Physiol 87:251–258.

Geran LC, Travers SP (2009) Bitter-responsive gustatory neurons in the rat parabrachial nucleus. J Neurophysiol 101:1598–1612.

Geran LC, Travers SP (2011) Glossopharyngeal nerve transection impairs unconditioned avoidance of diverse bitter stimuli in rats. Behav Neurosci 125:519–528.

Ginestal E, Matute C (1993) Gamma-aminobutyric acid-immunoreactive neurons in the rat trigeminal nuclei. Histochemistry 99:49–55.

Green DM, Swets JA (1966) Signal detection theory and psychophysics. New York,: Wiley.

Grossman SE, Fontanini A, Wieskopf JS, Katz DB (2008) Learning-related plasticity of temporal coding in simultaneously recorded amygdala-cortical ensembles. J Neurosci 28:2864–2873.

Gulbransen BD, Clapp TR, Finger TE, Kinnamon SC (2008) Nasal solitary chemoreceptor cell responses to bitter and trigeminal stimulants in vitro. J Neurophysiol 99:2929–2937.

Guo ZV, Li N, Huber D, Ophir E, Gutnisky D, Ting JT, Feng G, Svoboda K (2014) Flow of cortical activity underlying a tactile decision in mice. Neuron 81:179–194.

Halsell CB, Travers SP (1997) Anterior and posterior oral cavity responsive neurons are differentially distributed among parabrachial subnuclei in rat. J Neurophysiol 78:920–938.

Hamilton RB, Norgren R (1984) Central projections of gustatory nerves in the rat. J Comp Neurol 222:560–577.

Haring JH, Henderson TA, Jacquin MF (1990) Principalis-or parabrachial-projecting spinal trigeminal neurons do not stain for GABA or GAD. Somatosens Mot Res 7:391–397.

Herbert H, Moga MM, Saper CB (1990) Connections of the parabrachial nucleus with the nucleus of the solitary tract and the medullary reticular formation in the rat. J Comp Neurol 293:540–580.

Jordt SE, Bautista DM, Chuang HH, McKemy DD, Zygmunt PM, Hogestatt ED, Meng ID, Julius D (2004) Mustard oils and cannabinoids excite sensory nerve fibres through the TRP channel ANKTM1. Nature 427:260–265.

Kobayashi K, Fukuoka T, Obata K, Yamanaka H, Dai Y, Tokunaga A, Noguchi K (2005) Distinct expression of TRPM8, TRPA1, and TRPV1 mRNAs in rat primary afferent neurons with adelta/c-fibers and colocalization with trk receptors. J Comp Neurol 493:596–606.

Lemon C, Kang Y, Li J (2016) Separate functions for responses to oral temperature in thermo-gustatory and trigeminal neurons. Chem Senses 41:457–471.

Li J, Lemon C (2015a) Trigeminal convergence onto oral sensory neurons in the mouse nucleus of the solitary tract associates with gustatory and thermal tuning. Chem Senses 40:575.

Li J, Lemon CH (2015b) Influence of stimulus and oral adaptation temperature on gustatory responses in central taste-sensitive neurons. J Neurophysiol 113:2700–2712.

Liu L, Simon SA (1998) Responses of cultured rat trigeminal ganglion neurons to bitter tastants. Chem Senses 23:125–130.

Maier JX, Blankenship ML, Li JX, Katz DB (2015) A Multisensory Network for Olfactory Processing. Curr Biol 25:2642–2650.

Marfurt CF, Rajchert DM (1991) Trigeminal primary afferent projections to “non-trigeminal” areas of the rat central nervous system. J Comp Neurol 303:489–511.

McKemy DD, Neuhausser WM, Julius D (2002) Identification of a cold receptor reveals a general role for TRP channels in thermosensation. Nature 416:52–58.

Ninomiya Y, Sako N, Funakoshi M (1989) Strain differences in amiloride inhibition of NaCl responses in mice, Mus musculus. J Comp Physiol 166:1–5.

Ogura T, Krosnowski K, Zhang L, Bekkerman M, Lin W (2010) Chemoreception regulates chemical access to mouse vomeronasal organ: role of solitary chemosensory cells. PLoS One 5:e11924.

Ohkuri T, Horio N, Stratford JM, Finger TE, Ninomiya Y (2012) Residual chemoresponsiveness to acids in the superior laryngeal nerve in “taste-blind” (P2X2/P2X3 double-KO) mice. Chem Senses 37:523–532.

Oka Y, Butnaru M, von Buchholtz L, Ryba NJ, Zuker CS (2013) High salt recruits aversive taste pathways. Nature 494:472–475.

Otchy TM, Wolff SB, Rhee JY, Pehlevan C, Kawai R, Kempf A, Gobes SM, Olveczky BP (2015) Acute off-target effects of neural circuit manipulations. Nature 528:358–363.

Palmiter RD (2018) The Parabrachial Nucleus: CGRP Neurons Function as a General Alarm. Trends Neurosci 41:280–293.

Pittman DW, Contreras RJ (1998) Responses of single lingual nerve fibers to thermal and chemical stimulation. Brain Res 790:224–235.

Reed DR, Knaapila A (2010) Genetics of taste and smell: poisons and pleasures. Prog Mol Biol Transl Sci 94:213–240.

Ricardo JA, Koh ET (1978) Anatomical evidence of direct projections from the nucleus of the solitary tract to the hypothalamus, amygdala, and other forebrain structures in the rat. Brain Res 153:1–26.

Sagne C, El Mestikawy S, Isambert MF, Hamon M, Henry JP, Giros B, Gasnier B (1997) Cloning of a functional vesicular GABA and glycine transporter by screening of genome databases. FEBS Lett 417:177–183.

Samuelsen CL, Fontanini A (2017) Processing of Intraoral Olfactory and Gustatory Signals in the Gustatory Cortex of Awake Rats. J Neurosci 37:244–257.

Schwaber JS, Sternini C, Brecha NC, Rogers WT, Card JP (1988) Neurons containing calcitonin gene-related peptide in the parabrachial nucleus project to the central nucleus of the amygdala. J Comp Neurol 270:416–426, 398-419.

Scott TR, Mark GP (1987) The taste system encodes stimulus toxicity. Brain Res 414:197–203.

Simons CT, Boucher Y, Carstens E (2003a) Suppression of central taste transmission by oral capsaicin. J Neurosci 23:978–985.

Simons CT, Boucher Y, Carstens MI, Carstens E (2003b) Lack of quinine-evoked activity in rat trigeminal subnucleus caudalis. Chem Senses 28:253–259.

Sofroniew NJ, Vlasov YA, Andrew Hires S, Freeman J, Svoboda K (2015) Neural coding in barrel cortex during whisker-guided locomotion. Elife 4.

St. John SJ, Spector AC (1998) Behavioral discrimination between quinine and KCl is dependent on input from the seventh cranial nerve: implications for the functional roles of the gustatory nerves in rats. J Neurosci 18:4353–4362.

Tabachnick B, Fidell L (2001) Using Multivariate Statistics, 4 Edition. Boston: Allyn & Bacon.

Tizzano M, Cristofoletti M, Sbarbati A, Finger TE (2011) Expression of taste receptors in solitary chemosensory cells of rodent airways. BMC Pulm Med 11:3.

Tokita K, Boughter JD, Jr. (2016) Topographic organizations of taste-responsive neurons in the parabrachial nucleus of C57BL/6J mice: An electrophysiological mapping study. Neuroscience 316:151–166.

Veit L, Nieder A (2013) Abstract rule neurons in the endbrain support intelligent behaviour in corvid songbirds. Nat Commun 4:2878.

Wang Y, Erickson RP, Simon SA (1995) Modulation of rat chorda tympani nerve activity by lingual nerve stimulation. J Neurophysiol 73:1468–1483.

Wiegert JS, Mahn M, Prigge M, Printz Y, Yizhar O (2017) Silencing Neurons: Tools, Applications, and Experimental Constraints. Neuron 95:504–529.

Wilson DM, Lemon CH (2014) Temperature systematically modifies neural activity for sweet taste. J Neurophysiol 112:1667–1677.

Wilson DM, Boughter JD, Jr., Lemon CH (2012) Bitter taste stimuli induce differential neural codes in mouse brain. PLoS One 7:e41597.

Yamamoto T, Shimura T, Sakai N, Ozaki N (1994) Representation of hedonics and quality of taste stimuli in the parabrachial nucleus of the rat. Physiol Behav 56:1197–1202.

Yarmolinsky DA, Peng Y, Pogorzala LA, Rutlin M, Hoon MA, Zuker CS (2016) Coding and Plasticity in the Mammalian Thermosensory System. Neuron 92:1079–1092.

Zhao S, Ting JT, Atallah HE, Qiu L, Tan J, Gloss B, Augustine GJ, Deisseroth K, Luo M, Graybiel AM, Feng G (2011) Cell type-specific channelrhodopsin-2 transgenic mice for optogenetic dissection of neural circuitry function. Nat Methods 8:745–752.

Zotterman Y (1936) Specific action potentials in the lingual nerve of cat. Skand Arch Physiol 75:105–119.

